# Built on sand: the shaky foundations of simulating single-cell RNA sequencing data

**DOI:** 10.1101/2021.11.15.468676

**Authors:** Helena L. Crowell, Sarah X. Morillo Leonardo, Charlotte Soneson, Mark D. Robinson

## Abstract

With the emergence of hundreds of single-cell RNA-sequencing (scRNA-seq) datasets, the number of computational tools to analyse aspects of the generated data has grown rapidly. As a result, there is a recurring need to demonstrate whether newly developed methods are truly performant – on their own as well as in comparison to existing tools. Benchmark studies aim to consolidate the space of available methods for a given task, and often use simulated data that provide a ground truth for evaluations. Thus, demanding a high quality standard for synthetically generated data is critical to make simulation study results credible and transferable to real data.

Here, we evaluated methods for synthetic scRNA-seq data generation in their ability to mimic experimental data. Besides comparing gene- and cell-level quality control summaries in both one- and two-dimensional settings, we further quantified these at the batch- and cluster-level. Secondly, we investigate the effect of simulators on clustering and batch correction method comparisons, and, thirdly, which and to what extent quality control summaries can capture reference-simulation similarity.

Our results suggest that most simulators are unable to accommodate complex designs without introducing artificial effects; they yield over-optimistic performance of integration, and potentially unreliable ranking of clustering methods; and, it is generally unknown which summaries are important to ensure effective simulation-based method comparisons.

## Introduction

Single-cell RNA-sequencing (scRNA-seq) has become an established tool for studying the transcriptome at individual cell resolution. Since the first scRNA-seq study’s publication in 2009^[1]^, there has been a rapid increase in the number of scRNA-seq datasets, number of cells and samples per dataset^[2]^, and a corresponding growth in the number of computational methods to analyse such data, with over one thousand tools catalogued to date^[3,4]^. With the development of new methods comes the need to demonstrate their performance, and to consolidate the space of available methods through comprehensive and neutral benchmark studies^[5,6,7]^.

In this context, simulations have become an indispensable tool, for example, to investigate how methods respond to varying parameter inputs and quantify their scalability in terms of computational cost, as well as ensuring that a method is performant across a range of scenarios and in comparison to other available tools. The attractiveness of simulation studies is largely due to being able to specify a ground truth, which is often challenging or infeasible to establish in experimental data^[8]^. For example, evaluation of methods to group cells into biologically meaningful subpopulations (clusters) relies on ‘true’ labels to be compared against. While these may be attainable (e.g., through cell-sorting) or derived (e.g., manual annotation by an expert), simulations enable testing methods across a wide range of scenarios where the number of clusters, between-cluster (dis)similarity and effects of other covariates can be deeply explored. As a result, simulations have been applied to benchmark methods across a wide range of tasks, including differential expression analysis^[9,10,11]^, trajectory inference^[12]^, and data integration^[13,14]^.

By definition, simulations generate *synthetic* data. On the one hand, conclusions drawn from simulation studies are frequently criticized, because simulations cannot completely mimic (real) experimental data. On the other hand, it is often too expensive, or even impossible, to generate experimental data that is suitable for formal performance comparison. Nonetheless, setting a high quality standard for simulations is all the more important to ensure results based on them are transferable to corresponding experimental datasets.

Typically, new simulation methods come with minimal (non-neutral) benchmarks that focus on onedimensional evaluations, i.e. how similarly a set of summaries is distributed between a reference and simulated dataset (e.g., Zappia *et al*.^[15]^). In some cases, two-dimensional relationships (e.g., gene expression mean-variance) are explored (e.g., Assefa *et al*.^[16]^). However, the faithfulness of the full complexity of the simulated scRNA-seq data, including batch effects and clusters, is rarely evaluated. To date, there has been only one neutral evaluation of how well scRNA-seq data simulators recapitulate key characteristics of the counts, sample- and subpopulation-effects present in real data^[17]^; in particular, they proposed a novel kernel density metric to evaluate similarity of real and simulated data summaries. However, to what extent simulators affect the results of method comparisons is not considered.

Methods for simulating scRNA-seq data may be categorized according to various factors. Most importantly, there is a dichotomy between methods that generate synthetic data *de novo*, and those that rely on a reference dataset. The former depend on user-defined parameter inputs to generate counts, and introduce artificial effects between, e.g., different groups of cells or samples. Conversely, reference-based methods estimate parameters to mimic the gene expression profiles observed in the reference dataset. However, many methods employ a hybrid framework where, e.g., baseline parameters are estimated from a ‘singular’ reference (i.e., a homogeneous group of cells), and additional layers of complexity (e.g., batch effects, multiple clusters and/or experimental conditions) are added *post hoc*. Both strategies have their advantages and disadvantages: *de novo* simulators offer high flexibility in varying the strength and specificity of different effects, but might not generate realistic data; in contrast, reference-based methods are limited to the complexity of the input data and consequently less flexible, but are by default more realistic. Taken together, there is a trade-off between how applicable methods are in benchmarking single-cell analysis tools across a wide range of scenarios versus whether simulation study results are directly transferable to real data.

Here, we evaluated 16 scRNA-seq simulation methods in their ability to replicate important aspects of a real reference dataset. We considered various global, gene- and cell-level summaries, and compared them between reference and simulated data, in both one- and two-dimensional settings. In addition to global distributions (i.e. across all cells), we made batch- and cluster-level comparisons to capture structural differences in the summaries.

Our results suggest that there is a noticeable shortage of simulators that can accommodate complex situations, and that popular simulators do not adequately mimic real datasets. In particular, some current methods are able to simulate multiple groups of cells (e.g., batches and/or clusters), but do so in an *ad hoc* manner, e.g., by introducing arbitrary differences based on parameter inputs. Few methods attempt to estimate and mimic group effects from reference datasets, but this comes at a loss of supplying a ground truth.

## Results

### Benchmark design

We evaluated simulators based on 12 published datasets (see Reference datasets), from which we generated a variety of subsets that serve as references for simulation (Supp. Tab. 1). We labelled references as one of three types according to their complexity: type *n* are ‘singular’ references that contain cells from a single batch and cluster; type *b* contain cells from multiple batches; and type *k* contain cells from multiple clusters (Supp. Fig. 1-3). Here, batches can be either biological or technical replicates; clusters refer to cell subpopulations or types as annotated in the original data; and groups can be either batches, clusters, or experimental conditions. In total, we used 10, 8, and 8 references of type *n*, *b*, and *k*, respectively.

To objectively cover the space of currently available simulators, we browsed the scRNA-seq tools database^[3,4]^, which, at the time of writing, catalogued over 1000 tools for analysing scRNA-seq data, 65 of which hold a “simulation” tag. We included all methods that i) could be installed and run after at most minor manual adjustment(s); and, ii) were reference-based, i.e., supported parameter estimation from a real dataset. We selected a total of 16 methods, 9/6 of which could accommodate batches/clusters (Tab. 1). A brief summary of each method’s model framework, capabilities and limitations, as well as the parameter settings used in this study is given under Methods.

**Table 1:** Overview of scRNA-seq simulators compared in this study. Methods are ordered alphabetically and annotated according to their (in)ability to accommodate multiple batches and/or clusters, support for parallelization (parameter estimation and data simulation, respectively), software availability, and publication year. The right-most column catalogues neutral benchmark studies where each simulator was used. (✔ = yes, ✘ = no, (✔)= yes, but based on user input parameters, i.e. no support for parameter estimation, *requires random splitting of cells into two groups, †/‡ = internal/prior resampling from empirical parameter distribution, ◦ = no separate estimation step).

|  | Batches | Clusters | Type(s) | Cell # | Parallelization | Availability | Year | Model |
| --- | --- | --- | --- | --- | --- | --- | --- | --- |
| BASiCS<sup>[37]</sup> | ✓ | ✗ | b | ✗ | ✓ ✗ | R/Bioc | 2015 | NB |
| ESCO<sup>[38]</sup> | ✓ | ✓ | n,b,k | ✓ | ✓ ✓ | R/GitHub | 2020 | Gamma-Poisson |
| hierarchicell<sup>[39]</sup> | ✓ | ✗ | n,b | ✓ | ✗ ✗ | R/GitHub | 2021 | NB |
| muscat<sup>[40]</sup> | ✓ | ✓ | n,b,k | (✓)<sup>†</sup> | ✗ ✗ | R/Bioc | 2020 | NB |
| POWSC<sup>[41]</sup> | ✗ | ✓ | n,k | (✓)<sup>†</sup> | ✗ ✗ | R/Bioc | 2020 | zero-inflated, log-normal<br>Poisson mixture |
| powsimR<sup>[42]</sup> | ✗ | (✓) | n* | (✓)<sup>†</sup> | ✓ ✓ | R/GitHub | 2017 | NB |
| scDD<sup>[43]</sup> | ✗ | ✗ | n* | ✓ | ✓ ✓ | R/Bioc | 2016 | Bayesian NB mixture model |
| scDesign<sup>[44]</sup> | ✗ | (✓) | n | ✓ | ○ ✓ | R/GitHub | 2019 | Gamma-Normal mixture<br>model |
| scDesign2<sup>[45]</sup> | ✗ | ✓ | n,k | ✓ | ✓ ✗ | R/GitHub | 2020 | (zero-inflated) Poisson or NB<br>+ Gaussian copula for<br>gene-gene correlations |
| SCRIP<sup>[46]</sup> | ✓ | ✓ | n,b,k | ✓ | ✗ ✗ | R/GitHub | 2020 | (Beta-)Gamma-Poisson |
| SPARSim<sup>[47]</sup> | ✓ | ✗ | n,b | (✓)<sup>‡</sup> | ✗ ✗ | R/GitLab | 2020 | Gamma-multivariate<br>hypergeometric |
| splatter<sup>[15]</sup><br>(Splat model) | (✓) | (✓) | n | ✓ | ✗ ✗ | R/Bioc | 2017 | Gamma-Poisson |
| SPsimSeq<sup>[16]</sup> | ✓ | ✗ | n,b | ✓ | ○ ✗ | R/Bioc | 2020 | log-linear model-based density<br>estimation + Gaussian copula<br>for gene-gene correlations |
| SymSim<sup>[48]</sup> | ✓ | ✗ | n,b | ✓ | ✗ ✗ | R/GitHub | 2019 | kinetic model using MCMC |
| ZINB-WaVE<sup>[49]</sup> | ✓ | ✓ | n,b,k | ✗ | ✗ ✗ | R/Bioc | 2018 | zero-inflated NB |
| zingeR<sup>[50]</sup> | ✗ | ✗ | n | (✓)<sup>†‡</sup> | ✗ ✗ | R/GitHub | 2017 | zero-inflated NB |

Because different simulators can generate different levels of complexity (two groups, multiple clusters or batches, both or neither), we tagged each method according to their capabilities (type *n*, *b* and/or *k*). Each method was only run on corresponding reference datasets. A more detailed overview of the computational pipeline for this study is given in Fig. 1 (see also Computational workflow). Notably, there are various methods aimed at simulating continuous scenarios (e.g., a trajectory or time-course^[18,19,20]^). Here, we limited comparisons to methods that generate a single group or multiple groups of cells, since current continuous simulators are fully or in part *de novo*, making it challenging to validate the faithfulness of the data they generate (see Discussion).

**Figure 1:**
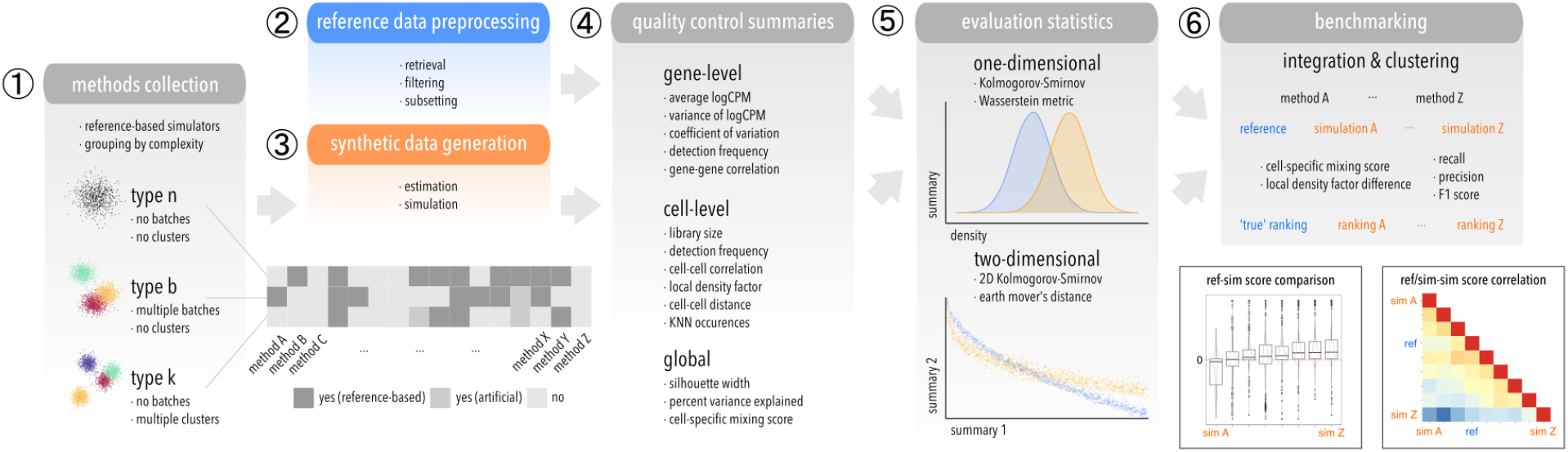
Schematic of the computational workflow used to benchmark scRNA-seq simulators. (1) Methods are grouped according to which level of complexity they can accommodate: type *n* (‘singular’), *b* (batches), *k* (clusters). (2) Raw datasets are retrieved reproducibly from a public source, filtered, and subsetted into various datasets that serve as reference for (3) parameter estimation and simulation. (4) Various gene-, cell-level and global summaries are computed from reference and simulated data, and (5) compared in a one- and two-dimensional setting using two statistics each. (6) Integration and clustering methods are applied to type *b* and *k* references and simulations, respectively, and relative performances compared between reference-simulation and simulation-simulation pairs.

In order to investigate how widely and for what purpose different simulators are applied, we browsed the literature for benchmark studies that compare tools for a specific scRNA-seq analysis task. Depending on the task, such comparisons often rely on simulation studies where a ground truth is known (by design), or a combination of simulated and real data, where an experimental ground truth exists. For each benchmark, we summarized the task of interest and, if any, which simulator(s) are used. Across all considered benchmarks, these amounted to only five, namely: `muscat` (1^[21]^), `scDesign` (1^[22]^), `powsimR` (2^[10,23]^), `scDD` (3^[9,11,24]^), and `splatter` (13^[12,13,14,25,26,27,28,29,30,31,32,33,34]^). Benchmark tasks included batch effects^[13,14]^, clustering^[28,31,35]^, doublet detection^[22]^, differential expression^[9,10,11,30]^, dimensionality reduction^[29,32]^, imputation^[25,34]^, isoform quantification^[36]^, marker selection^[27,24]^, normalization^[26]^, pipelines^[23,21]^, cell type assignment^[33]^, and trajectory inference^[12]^. Yet, this listing of benchmarks and their use cases is not exhaustive; the frequency with which simulators are applied in benchmarks need not speak for or against their performance (e.g., long-lived and user-friendly methods might be favored); and, because some simulators have not been applied to a given task does not mean they can not be.

In order to summarize how well each simulator recapitulates key characteristics of the reference scRNA-seq dataset, we computed a range of gene- and cell-level summaries for both reference and simulated datasets. These include average and variance of log-transformed counts per million (CPM), coefficient of variation, gene detection frequency, gene-to-gene correlation, log-transformed library size (total counts), cell detection frequency, cell-to-cell correlation, local density factor, cell-to-cell distance, and k-nearest neighbor (KNN) occurrences (see Quality control summaries).

Since some summaries (e.g., detection frequency) can vary between batches (e.g., sequencing depths may vary between protocols) and clusters (e.g., different cell types may differ in their overall expression), we computed them globally, i.e. across all cells, as well as for each batch and cluster. Thus, for a given dataset with *B* batches and *K* clusters, we obtain 1, 1 + *B*, and 1 + *K* results per summary for type *n*, *b*, and *k*, respectively. Three additional summaries were computed across all cells – namely, the percent variance explained (PVE) at the gene-level (i.e. expression variance accounted for by batch/cluster for type *b/k*); and the silhouette width and cell-specific mixing score (CMS) at the cell-level (considering as group labels the batch/cluster for type *b/k*) – that aim to capture global batch or cluster effects on gene expression variability and cell-to-cell similarity, respectively.

To evaluate simulator performance, we compared summaries between reference and simulated data in one- and two-dimensional settings by computing the Kolmogorov-Smirnov (KS) distance^[51]^ and Wasserstein metric for each summary (Supp. Fig. 5-7), and the KS distance and earth mover’s distance (EMD) for each relevant pair of summaries (Supp. Fig. 8-10). In general, these metrics quantify how dissimilar a pair of (univariate or bivariate) distributions are (see Evaluation statistics). Test statistics were generally consistent between KS test and Wasserstein metric, as well as KS test and EMD (Supp. Fig. 4). Thus for brevity, method performances are hereafter reported as one- and two-dimensional KS statistics.

### Simulators vary in their ability to mimic scRNA-seq data characteristics

Across all simulation types, cell-level quality control summaries were generally poorly recapitulated (Fig. 2a), with the largest deviation in cell-to-cell correlation. The silhouette width, CMS and PVE gave amongst the highest KS distances for most methods, indicating that, while group-level (i.e., within a batch or cluster) summaries might be preserved well during simulation, the global data structure (e.g., inter-group relations) is not. Despite its popularity, `splatter` ranked in the middle for the majority of summaries. `scDD` ranked poorly for most summaries, preceded by `hierarchicell` and `ESCO`. Considering all summaries, `ZINB-WaVE`, `scDesign2`, and `muscat` were among the best performing simulators, yielding low KS test statistics across a large number of metrics and datasets (Fig. 2b).

**Figure 2:**
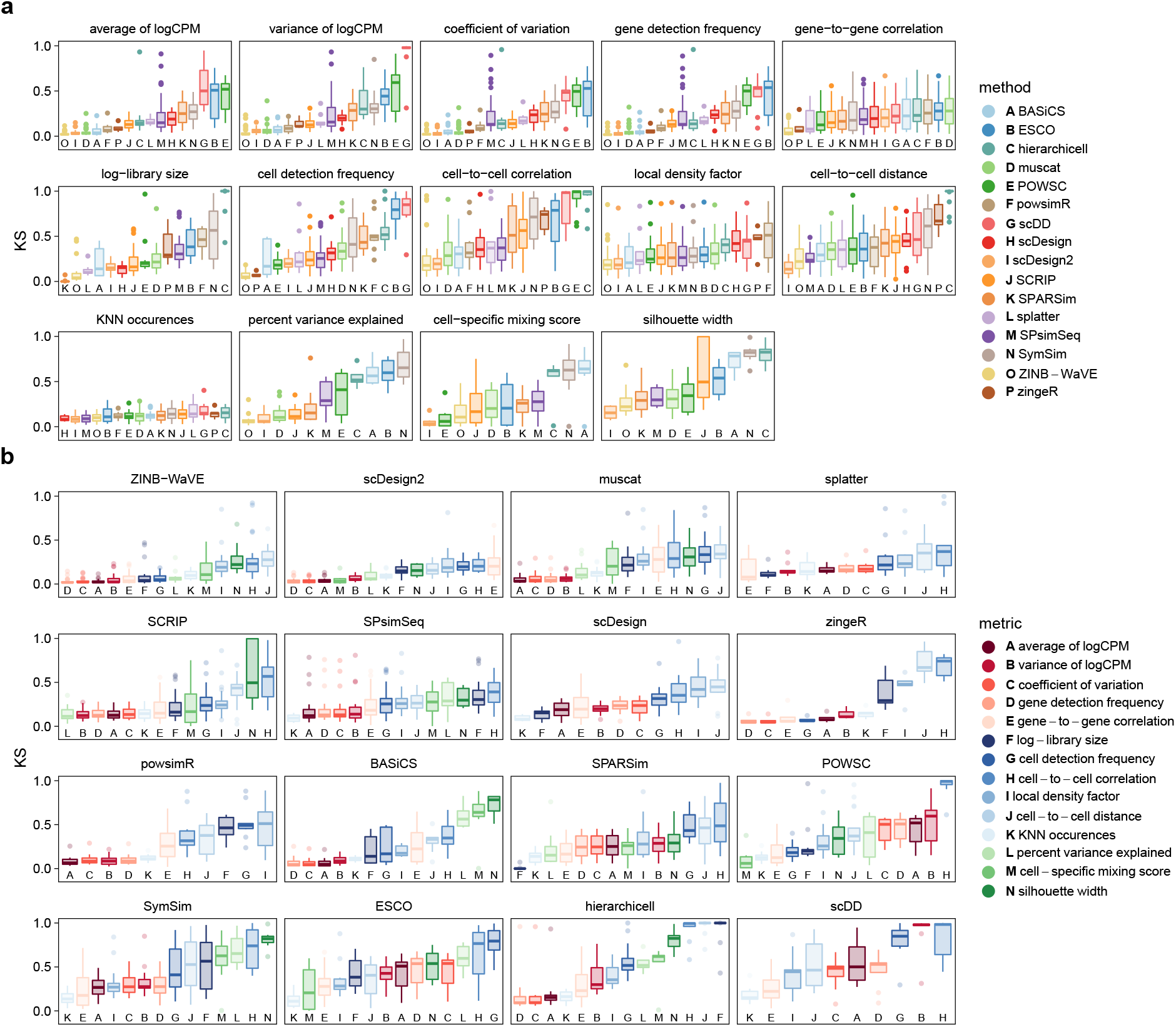
Kolmogorov-Smirnov (KS) test statistics comparing reference and simulated data across methods and summaries. Included are datasets and methods of all types; statistics are from global comparisons for type *n*, and otherwise averaged across cluster-/batch-level results. (a) Data are colored by method, and stratified by summary. For each summary (panel), methods (x-axis) are ordered according to their average. (b) Data are colored by summary, and stratified by method. For each method (panel), metrics (x-axis) are ordered according to their average from best (small) to worst (large KS statistic). Panels (methods) are ordered by increasing average across all summaries.

Finally, we ranked simulators according to their overall performance. In order to weight datasets equally and independently of the number of subsets drawn from them, we first averaged statistics across subsets, then datasets. Secondly, because simulators ranked similarly in both one- and two-dimensional comparisons, and performances were often linked for certain metrics, we limited rankings to one-dimensional evaluations only, and averaged across all gene- and cell-level metrics. This resulted in three independent rankings, one for each set of methods that can accommodate a given simulation type (Fig. 3). Notably, subpopulations (batches/clusters) may vary in size and complexity. Thus, for types other than *n*, we averaged across group-level results (instead of using global test results).

**Figure 3:**
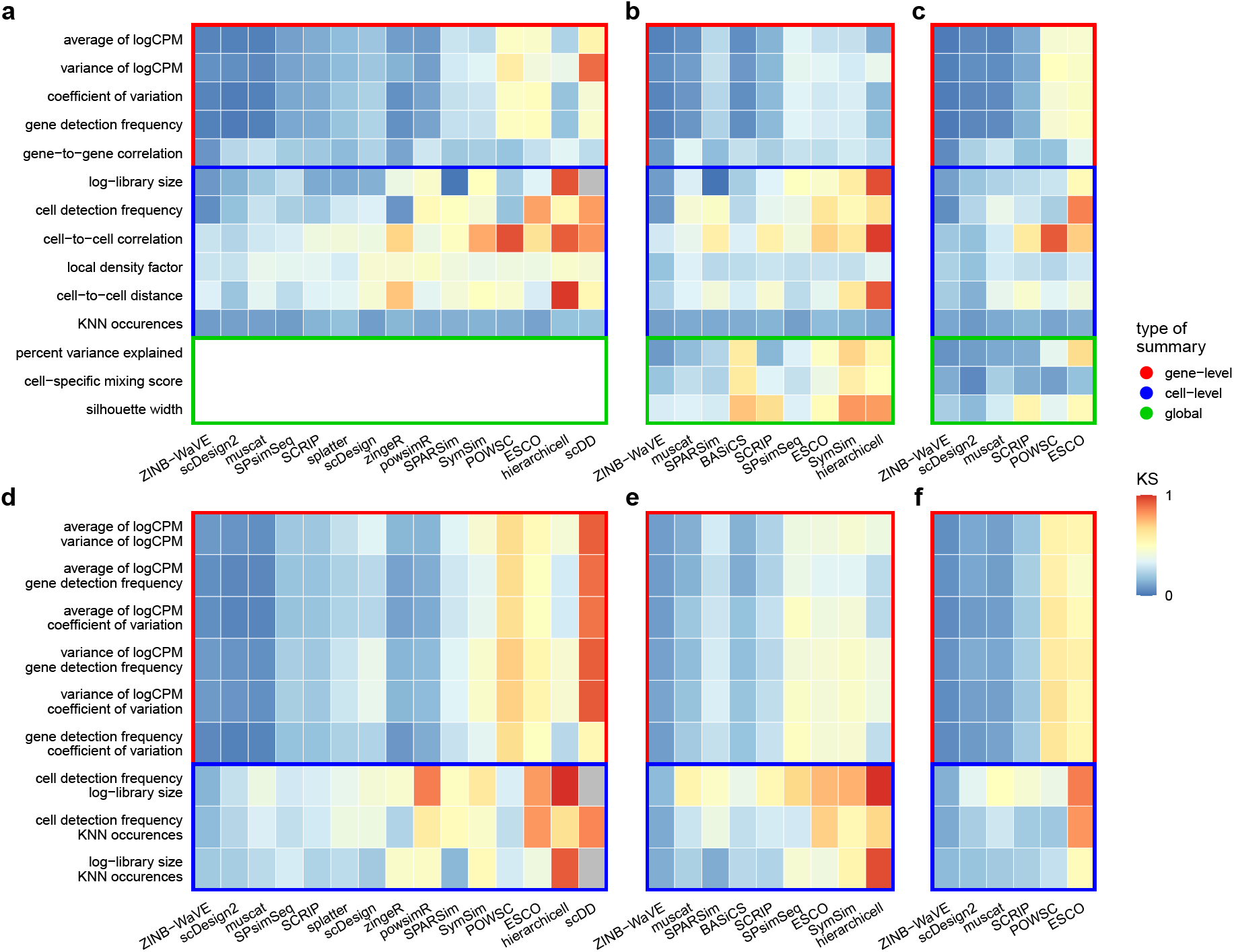
Average performance in one-(upper row) and two-dimensional evaluations (bottom row) for (a,d) type *n*, (b,e) type *b*, and (c,f) type *k* simulations. For each type, methods (x-axis) are ordered according to their average performance across summaries in one-dimensional comparisons. Except for type *n*, batch- and cluster-level results are averaged across batches and clusters, respectively. Boxes highlight gene-level (red), cell-level (blue), and global summaries (green).

For type *n*, `ZINB-WaVE`, `scDesign2`, `muscat`, and `SPsimSeq` performed similarly well, with `POWSC`, `ESCO`, `hierarchicell`, and `scDD` ranking last across various summaries. `ZINB-WaVE` and `muscat` were also the most performant among type *b* and *k* simulators, joined by `SPARSim` and `scDesign2`, respectively. LDF, cell-to-cell distance and correlation (across all types) and global summaries (PVE and silhouette width for type *b* and *k*) were poorly recapitulated.

To measure the scalability of methods, we repeatedly timed estimation and simulation steps across varying numbers of genes and cells (see Methods). Runtimes varied across several orders of magnitude (Supp. Fig. 11). Some methods did not offer separate estimation and data generation steps, while others can generate multiple simulations from one-time parameter estimates. Specifically, the estimation step of `scDesign2`, `ZINB-WaVE` and `zingeR` was relatively slow, but data simulation was not. In contrast, `scDD` and `SymSim` took longer for simulation than estimation. Overall, `BASiCS` was by far the slowest. `SPsimSeq`, `SPARSim`, `SCRIP` and `SymSim` were approximately ten fold faster. The remaining methods (`ESCO`, `hierarchicell`, `muscat`, `POWSC`, and `splatter`) were the fastest. Memory usage (Supp. Fig. 12 and Supp. Fig. 13) was similar across methods but exceptionally high for `SPSimSeq`. While some methods provide arguments for parallelization (see Tab. 1), all methods were run on a single core for comparability.

### Batch simulators yield over-optimistic but faithful integration method performance

Ideally, benchmark results (i.e. the ranking of computational tools for a given task) should be the same for experimental and simulated data. In order to investigate how method comparison results are affected by simulation, we used the 8 type *b* references to compare 6 scRNA-seq batch correction methods. To evaluate method performances, we computed: i) cell-specific mixing scores (CMS); and, ii) difference in local density factors (ΔLDF) before and after integration^[52]^. In order to make metrics comparable across datasets, we zero-centered CMS (denoted CMS*), and zero-centered and range-one scaled ΔLDF (denoted ΔLDF*). Finally, we computed a batch correction score BCS = |CMS*| + |ΔLDF*|, where small values indicates ideal mixing (CMS* of 0 on average) while retaining the data’s internal structure (ΔLDF* centered at 0), and large values indicates batch-specific bias (high CMS* density at ±1) and changes in overall structure (ΔLDF* non-symmetric).

ΔLDF* were largely consistent between references and simulations (Supp. Fig. 14), whereas CMS* were much less correlated for most methods (Supp. Fig. 15). BCSs were overall similar for simulated compared to reference data, and well correlated between most reference-simulation and simulation-simulation pairs (Supp. Fig. 16). Simulations from `SPsimSeq`, `ZINB-WaVE`, `SPARsim` and `SCRIP` gave results most similar to real data, followed by `BASiCS`, and lastly `muscat` and `SymSim` (see also Supp. Fig. 17-24).

### Cluster simulators affect the performance of clustering methods

Secondly, we used the 8 type *k* references to evaluate 9 scRNA-seq clustering methods that were previously compared in Duò *et al*.^[31]^. To evaluate method performances, we computed cluster-level F1 scores, after using the Hungarian algorithm^[53]^ to match cluster assignments to ‘true’ labels.

Across all methods and datasets, F1 scores were consistently higher for simulated compared to real data (Fig. 4a-b). In addition, for similarly performant simulators, clustering methods rankings were more dependent on the underlying reference dataset than the specific simulator used (Fig. 4c). And, some simulators (e.g., `SCRIP`) gave almost identical F1 scores and rankings for multiple references. Overall, method rankings (according to F1 scores) were lowly correlated between simulated and reference data, as well as between simulations (Fig. 4d), with `scDesign2` and `POWSC` giving the most, and `muscat` and `SCRIP` giving the least similar ranking, respectively.

**Figure 4:**
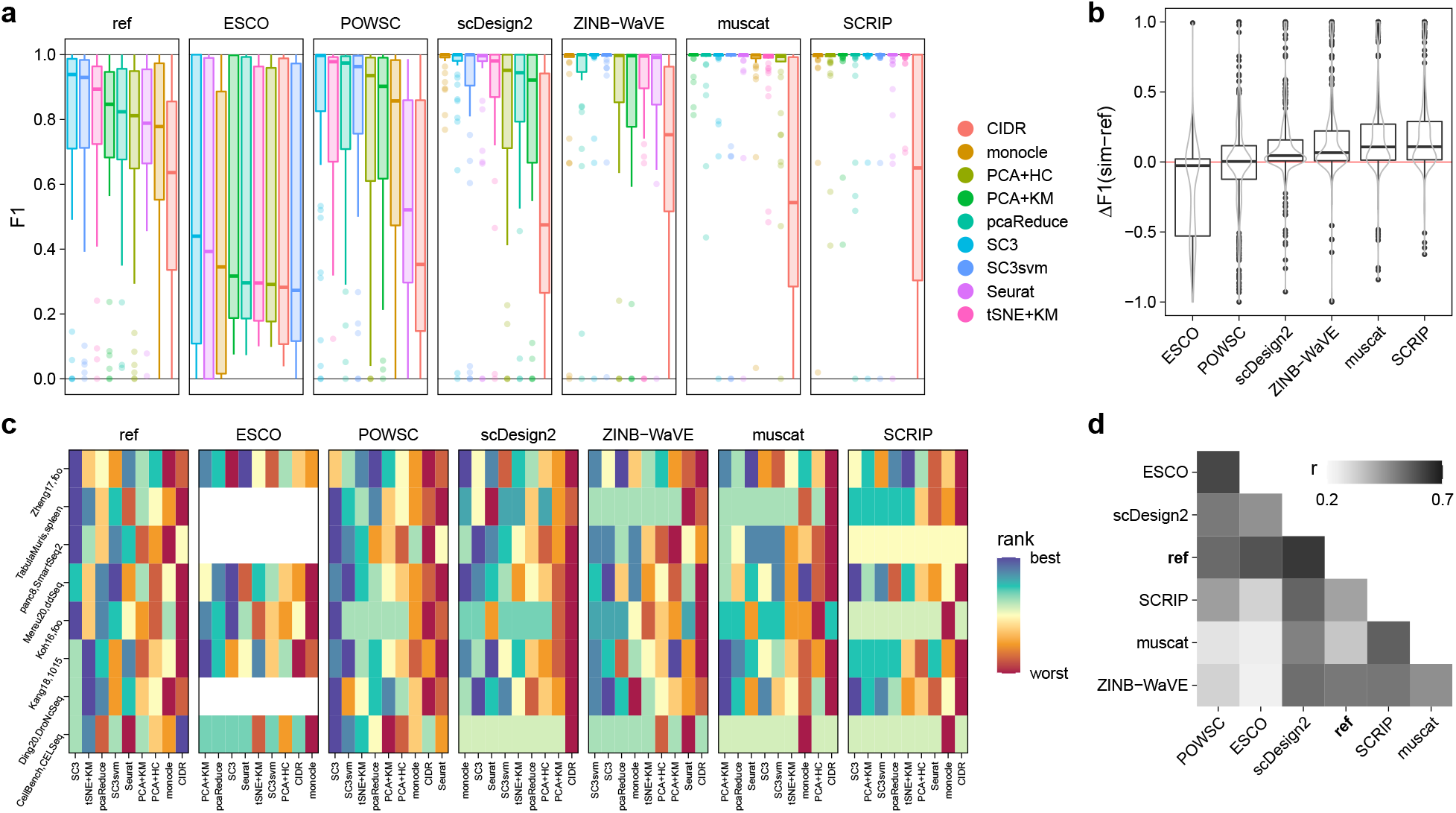
Comparison of clustering results across (experimental) reference and (synthetic) simulated data. (a) Boxplot of F1 scores across all type *k* references, simulation and clustering methods. (b) Boxplot of difference (Δ) in F1 scores obtained from *ref*erence and *sim*ulated data. (c) Heatmap of clustering method (columns) rankings across datasets (rows), stratified by simulator (panels). (d) Heatmap of Spearman’s rank correlation (*ρ*) between F1 scores across datasets and clustering methods.

Taken together, these results suggest that simulations do not achieve the same level of complexity in terms of intra- and inter-subpopulation effects (i.e., batches and clusters). Consequently, methods to correct of such effects (integration) or group together similar cells (clustering) perform over-optimistically in simulated data compared to more complex and noisy experimental data, and are almost indistinguishable in their performance for ‘simple’ datasets.

### Meta-analysis of summaries

Inevitably, summaries used to assess whether simulated data mimics real data may be redundant, and we expect that some summaries are more important than others. To quantify the relationship between summaries, we correlated all comparable summaries, i.e., gene- and cell-level summaries, respectively, excluding those that include sampling, i.e., correlations and cell-to-cell distances (Fig. 5a). Gene detection frequency and average expression were highly similar (*r* ~ 1), and correlated well with expression variance (*r* > 0.5). At the cell level, detection frequencies and library sizes were most similar.

**Figure 5:**
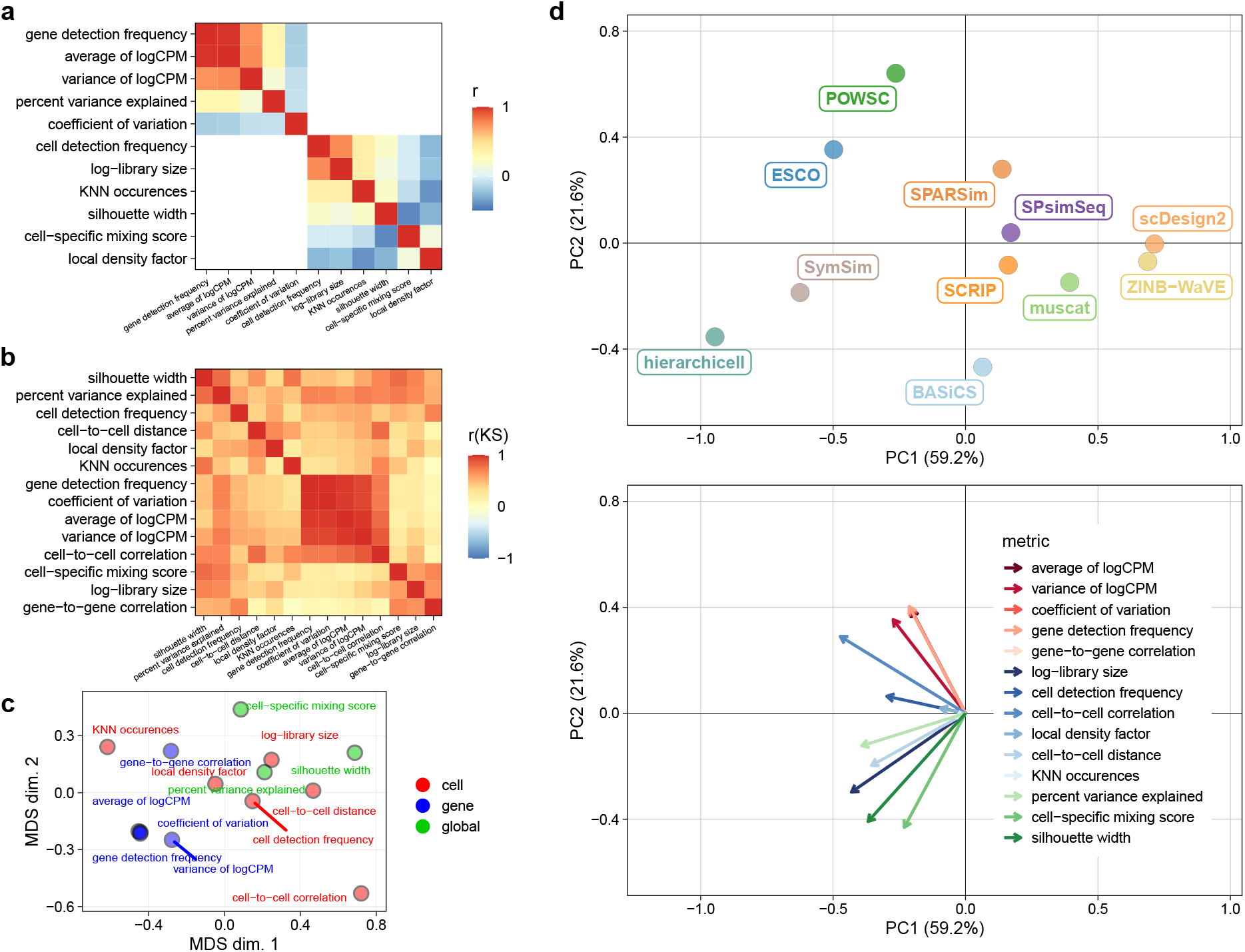
Comparison of quality control summaries and KS statistics across datasets and methods. Spearman rank correlations (r) of (a) gene- and cell-level summaries across reference datasets, and (b) KS statistics across methods and datasets. (c) Multi-dimensional scaling (MDS) plot, and (d) principal component (PC) analysis of KS statistics across all and type *b/k* methods, respectively, averaged across datasets.

Next, we correlated the KS test statistics obtained from comparing reference-simulation pairs of summaries across all datasets (Fig. 5b). Summaries grouped together according to their type (global, gene- or cell-level), indicating that simulators recapitulated each type of summary to a similar degree, and that one summary per type could be sufficient to distinguish between method performances.

To investigate the overall similarity of summaries, we performed multi-dimensional scaling (MDS) on KS statistics across methods and datasets (Fig. 5c and Supp. Fig. 25). In line with the observed correlation structure, summaries grouped together by type, with gene-level summaries being most similar to one another.

Next, we performed principal component analysis (PCA) on test statistics of summaries across methods and datasets (Fig. 5d and Supp. Fig. 26-28), thus representing each method-dataset as a linear combination of statistics for each summary. For all types, the largest fraction of variance (PC1: >40%) was attributable to differences in overall method performance, followed by (PC2: >15%) differences in summary type-specific performance (global, gene-, cell-level).

Taken together, our analyses suggest that several summaries convey similar information, with gene-level summaries being particularly redundant, and global summaries the least redundant. Accordingly, simulator performance may be sufficiently quantifiable by a combination of one gene-level, few cell-level, and various global summaries. However, the number (and nature) of summaries to comprehensively cover the inherent data structure is dependent on its complexity (e.g., global summaries are void for type *n*, but all the more relevant for type *b* and *k*); and there may exist other informative summaries not considered here.

## Discussion

In this study, we compared scRNA-seq data simulators in their ability to generate synthetic data that can recapitulate key characteristics of real reference datasets. We considered a range of gene- and cell-level summaries, as well as ones specific to capturing local and global group-effects (i.e. intra- and inter-variability of batches and clusters). By comparing the distribution of summaries as well as pairs thereof between reference and simulated data, we evaluated how well simulations capture the structure exhibited by a given reference dataset. We ranked simulators by averaging their performance across summaries and simulations, thus evaluating each method across a multifaceted range of criteria (covering gene-, cell- and group-specific summaries) and datasets (from various tissues and technologies). Our results confirm some aspects of those in the study from Cao *et al*.^[17]^ (our study includes 16 simulators, theirs covered 12, with an overlap of 10); for example, `ZINB-WaVE` stands out as performant in nearly all reference-simulation comparisons. However, we additionally investigated whether the choice of simulator affects method comparison results, included additional quality control summaries that capture group (batch/cluster) structure, and stratified simulators according to type (singular, batch, cluster).

Overall, simulations have proven paramount for the development of computational tools for the analysis of single-cell data. However, they are often implemented for a specific task, e.g., evaluating clustering, batch correction, or DE analysis methods; an ‘all-rounder’ simulator is currently lacking, and most methods are limited to simple designs. The arguably most sobering observation of this study is that, without introducing arbitrary effects that depend on user inputs (e.g., the frequency of DE genes and magnitude of changes in their expression), the vast majority of methods can simulate only one group of cells (i.e., one batch, one cluster). As a result, most current simulators are rather limited in their applicability.

Both neutral benchmark studies as well as non-neutral comparisons performed alongside newly presented methods rely on a ground truth for evaluation. Thus, future work should be focused on the development of flexible, faithful simulation frameworks to fill this gap, especially in scenarios where an experimental ground truth is challenging or infeasible to establish. For example, which genes and cell subpopulations are affected by batch effects cannot be controlled, independent of whether control samples might be used to quantify these effects. Similarly, intra- and inter-cluster effects are unclear, even if cluster annotations might be obtained through cell-sorting or manual annotation by an expert. And, effects on gene expression remain unknown, despite controlled perturbation or time-series studies through which discrete labels might be given. Taken together, although some level of ground truth may be experimentally attainable, simulations remain indispensable owing to: i) their feasibility; and, ii) the information they provide (e.g., which genes and cell subpopulations are affected).

The most truthful model for real data is real data. Artificial data alterations (e.g., applying fold changes to a specified subset of gene expression means in certain subsets of cells) are unlikely to mimic biological differences. Even if founded on a thorough investigation of *realistic* changes, non-reference based simulations are difficult to evaluate, and conclusions drawn from *de novo* simulations in terms of method evaluations should be treated with caution.

While tools to evaluate the quality of simulated data exist, they are seldomly taken advantage of. For example, `scater`^[54]^ offers a range of gene- and cell-level quality control summaries; `countsimQC`^[55]^ can generate a comprehensive report comparing an input set of count matrices (e.g., real against synthetic data); and, many dataset summaries are easy to compute and compare manually. Having such reference-simulation comparisons available every time that a simulator is proposed or used, as well as in every (non-neutral) benchmark would add credibility to the results.

In addition to evaluating the faithfulness of simulated data, we investigated whether and to what extent benchmark results are affected by the simulator used. Our results suggest that method performances for integration and clustering of scRNA-seq data deviate from those obtained from real data; in addition, simulators that better mimic reference datasets do not necessarily yield more similar method comparison results. For example, `muscat` was among the highest ranked simulators in our study, but integration and clustering method ranking obtained from `muscat` simulations were rather inconsistent with those from real data. On the other hand, `SPsimSeq` ranked mediocre in terms of mimicking real datasets, but gave the most faithful integration method ranking. In the context of clustering, there was a consistent over-optimistic performance of methods, independent of the simulator used.

This discrepancy between the faithfulness of simulated data and benchmark results brings to question which set of summaries is sufficient to capture relevant data structure. Here, simulators were ranked by their average performance across summaries. However, many of these may be redundant (see below), or differ in their suitability to capture group-related structures (e.g., batch-/cluster-effects). Thus, simulators that are performant ‘overall’ are not guaranteed to be suitable for evaluating methods for a specific task (e.g., integration/clustering), where global structure should take priority over gene-/cell-specific summaries. An open question that needs to be answered is what summaries are important for a given task.

Besides the capabilities each method has to offer and its performance, i.e. how realistic its simulations are, there are other criteria we did not explore thoroughly. For example, `splatter` offers a well-documented, easy-to-use framework that is both flexible and interpretable. While other methods might outperform `splatter`, they return parameters that are less applicable to benchmarking computational tools. For example, artificially introducing DE genes provides a binary ground truth (e.g., whether a gene is DE), whereas estimating and mimicking cluster effects might not (i.e., the user defines which genes are DE based on gene-wise parameters returned by the simulator).

Here, we have focused on methods that generate a single group or multiple groups of cells; in particular, we distinguished between ‘singular’ (type *n*), multi-batch (type *b*), and multi-cluster (type *k*) datasets. However, there are various methods that are aimed at simulating data where gene expression profiles evolve along a discrete or continuous trajectory or time-course (e.g., `dyngen`^[18]^, `PROSSTT`^[19]^, `SERGIO`^[20]^). These have been applied in, for example, benchmarking methods for trajectory inference^[12]^.

The combination of scRNA-seq with CRISPR/Cas9 genome editing has enabled the joint readout of gene expression and cell lineage barcodes^[56]^. Salvador-Martínez *et al*.^[57]^ have proposed a simulator for lineage barcode data that, however, does not generate gene expression data. A recent method, `TedSim`^[58]^, is capable of outputting combined readouts, and can be used to study tools for either or both data types, including more genuine investigation of trajectory inference methods.

Most of these methods employ fairly sophisticated and well-designed models for data generation, but require a complex set of inputs that is specific to each method and difficult to justify. Meanwhile, very few trajectory simulators support the estimation of simulation parameters from a reference dataset, making it challenging to evaluate them and opening the question of how faithful performance assessments based on them are. Overall, validating the faithfulness of synthetically generated trajectories in single-cell data remains challenging.

Taken together, while a set of performant methods to generate synthetic scRNA-seq data exist, current methods are: i) limited in the level of complexity they are able to accommodate; ii) often reliant - in full or in part - on inputs by the user to introduce (artificial) expression differences; iii) more or less suitable to evaluate other tools, depending on the data characteristics they can capture faithfully. Secondly, simulation-based benchmark studies are affected by the simulator used, and more performant simulators do not necessarily yield more reliable readouts of, e.g., integration and clustering methods. And thirdly, the chosen quality control summaries and their prioritization have an impact on the assessment of simulations and, consequently, the conclusions drawn from them. Thus, identifying the nature, number, and significance of summaries to faithfully capture scRNA-seq data structure warrants future work in order to improve method evaluations.

## Methods

### Reference datasets

Each reference dataset was retrieved from a publicly available source, including Bioconductor’s `ExperimentHub`^[59]^, the Gene Expression Omnibus (GEO) database, and public GitHub repositories. Raw data were formatted into objects of class `SingleCellExperiment`^[60,61]^ and, with few exceptions, left *as is* otherwise. Datasets cover various organisms, tissue types, technologies, and levels of complexity (i.e. number of genes and cells, clusters and/or batches and/or experimental conditions). A summary of each dataset’s characteristics and source is given in Supp. Tab. 1.

References underwent minimal filtering in order to remove groups (clusters, batches) with an insufficient number of cells, as well as genes and cells of low quality (e.g., low detection rate, few counts overall). Secondly, we drew various subsets from each reference to retain a reduced number of observations (genes and cells), as well as a known number of batches, clusters or neither (see Supp. Tab. 2 and Preprocessing).

### Simulation methods

With few exceptions, methods were run using default parameters and, if available, following recommendations given by the authors in the corresponding software documentation. All packages were available from a public GitHub repository, through CRAN, or Bioconductor^[62]^. A brief overview of each method’s model framework and support for parallelization is given in Tab. 1. For the explicit arguments used for parameter estimation and data simulation, we refer to the method wrappers available at https://github.com/HelenaLC/simulation-comparison (snapshot at DOI:10.5281/zenodo.5678950).

### Quality control summaries

We computed a set of five summaries at the gene-level: average and variance of logCPM, coefficient of variation, detection frequency (i.e. proportion of cells with non-zero count), as well as gene-to-gene correlation of logCPM. Here, logCPM correspond to log1p-transformed counts per million computed with `scater`’s `calculateCPM` function^[54]^. We also computed six summaries at the cell-level: library size (i.e. total counts), detection frequency (i.e. fraction of detected genes), cell-to-cell correlation (of logCPM), cell-to-cell distance (Euclidean, in PCA space), the number of times a cell occurs as a k-nearest neighbor (KNN), and local density factors^[52]^ (LDF) that represent a relative measure of a cell’s local density compared to those in its neighbourhood (in PCA space), and aim to quantify group (batch/cluster for type *b/k*) structure. For datasets other than type *n*, each summary was computed for each of three cell groupings: globally (i.e. across all cells), at the batch-, and at the cluster-level. Three additional summaries – the percent variance explained^[63]^ (PVE) at the gene-, and the cell-specific mixing score^[52]^ (CMS) and silhouette width^[64]^ at the cell-level – were computed globally. Here, the PVE corresponds to the fraction of expression variance accounted for by a cell’s group assignment (batch/cluster for type *b/k*). Summaries are described in more detail in Supp. Tab. 3.

### Evaluation statistics

For each reference-simulation pair of summaries, we computed the Kolmogorov-Smirnov (KS) test statistic using the `ks.test` function of the `stats` R package, and the Wasserstein metric using the `wasserstein_metric` function of the `waddR` R package^[65]^. In addition, we computed the two-dimensional KS statistic^[66]^ and earth mover’s distance (EMD) between relevant pairs of summaries, i.e. between unique combinations of gene- and cell-level summaries, respectively, excluding global summaries (PVE, CMS and silhouette width) as well as gene-to-gene and cell-to-cell correlations. One- and two-dimensional evaluations are detailed under Evaluation statistics.

### Runtime evaluation

To quantify simulator runtimes, we selected one reference per type, and drew five random subsets of 400-4,000 genes (fixing the number of cells) and 100-2,600 cells (fixing the number of genes). For each method and subset (eight in total), we separately measured the time required for parameter estimation and data simulation. For each step, we set a time limit of 10^6^ seconds after which computations were interrupted.

### Integration evaluation

Integration methods were implemented as in Chazarra-Gil *et al*.^[67]^ (see Integration). To evaluate method performances, cell-specific mixing scores (CMS) and the difference in local density factors (ΔLDF) were computed using the `cms` and `ldfDiff` function, respectively, of the `CellMixS` package^[52]^. To make metrics more interpretable and comparable across datasets, we i) subtracted 0.5 to center CMS at 0 (denoted CMS*); and, ii) centered (at 0) and scaled (to range 1) ΔLDF (denoted ΔLDF*). Overall integration scores correspond to the unweighted average of CMS* and ΔLDF*. Thus, for all three metrics, a value of 0 indicates ‘good’ mixing for a given cell. When aggregating results (e.g., for heatmap visualizations), metrics were first averaged across cells within each batch and, secondly, across batches.

### Clustering evaluation

Clustering methods were implemented as in Duò *et al*.^[31]^ (see Clustering). If applicable, the number of clusters was set to match the number of true (annotated respective simulated) clusters. To evaluate the performance of each method, we matched true and predicted cluster labels using the Hungarian algorithm, and computed cluster-level precision, recall, and F1 score (the harmonic mean of precision and recall).

## Data availability

All reference scRNA-seq datasets are available through Bioconductor’s `ExperimentHub`^[59]^, the Gene Expression Omnibus (GEO) database, or public GitHub repositories; see Supp. Tab. 1 for dataset-specific sources. R objects (*.rds* files) to reproduce key results of this study are available at DOI:10.5281/zenodo.5678871. These include global, gene- and cell-level quality control summaries of reference and simulated data for different cell groupings, one- and two-dimensional test statistics across all datasets and methods, clustering and integration results for reference and simulated data, and runtimes for 5 replicates per gene- and cell-subsets for one dataset per type; see Supplementary data for a comprehensive description.

## Code availability

All analyses were run in R v4.1.0^[68]^, with Bioconductor v3.13^[62]^. The computational workflow was implemented using `Snakemake` v5.5.0^[69]^, with Python v3.6.8. Package versions used throughout this study are captured in the *session_info.txt* file at DOI:10.5281/zenodo.5678871. All code to reproduce the results presented herein is accessible at https://github.com/HelenaLC/simulation-comparison (snapshot at DOI:10.5281/zenodo.5678950). Workflow structure, code organization and script naming schemes are described in more detail on the repository’s landing page.

## Author contributions

HLC implemented method comparisons with significant contributions from SM. CS assisted in several conceptual aspects of the benchmark. HLC, SM and MDR drafted the manuscript with feedback from CS. All authors read and approved the final paper.

## Acknowledgements

The authors thank members of the Robinson Lab at the University of Zurich for valuable feedback on methodology, benchmarking, and exposition. This work was supported by the Swiss National Science Foundation (grant Nos. 310030_175841 and CRSII5_177208) and the Chan Zuckerberg Initiative DAF, an advised fund of Silicon Valley Community Foundation (grant No. 2018-182828). MDR acknowledges support from the University Research Priority Program Evolution in Action at the University of Zurich.

