## Supplementary Text for "Built on sand: the shaky foundations of simulating single-cell RNA sequencing data"

### Contents

**1 Reference datasets** **1**

**2 Quality control summaries** **3**

**3 Method parameters** **4**

**4 Evaluation statistics** **5**

### 1 Reference datasets

Supp. Tab. 1 briefly summarizes each reference dataset, including the platform(s) on which scRNA-seq measurements were obtained, organism and tissue type, as well as the dataset's size (number of genes and cells), complexity (e.g. how many clusters, batches, whether batches are biological or technical replicates etc.) and (if applicable) how data were filtered or preprocessed. For more details, we refer readers to the associated publication(s). The subset(s) drawn from each dataset are summarized in Supp. Tab. 2; these serve as references for simulation.

| Dataset | Description | Preprocessing | Batches | Clusters | Features | Observations | Source |
|---|---|---|---|---|---|---|---|
| CellBench[1] | three human lung adenocarcinoma cell lines (HCC827, H1975, H2228) mixed in equal proportions and sequenced across three different platforms (CEL-Seq2, Drop-Seq, Chromium) | – | 3 | 3 | 13575 | 1401 | GSE118767 |
| Gierahn17[2] | human HEK293 (embryonic kidney cells) cell line sequenced with Seq-Well | – | – | – | 24187 | 1453 | GSE92495 |
| Ding20[3] | two mouse cortex snRNA-seq experiments (Cortex1 and Cortex2), each comprising 4 technologies (10x Chromium, DroNc-seq, sci-RNA-seq, Smart-Seq2) | retaining only first experiment (Cortex1) and cells that received a type annotation | 4 | 8 | 28692 | 4523 | SCP425 |
| Kang18[4] | droplet-based scRNA-seq data of PBMCs from eight patients, each measured before and after 6h treatment with IFN-$\beta$ | retaining untreated samples only, removing multiplets and cells that did not receive a type annotation | 8 | 8 | 17198 | 12315 | GSE96583 |
| Koh16[5] | in vitro cultured H7 human embryonic stem cells (WiCell) and H7-derived downstream early mesoderm progenitors | – | – | 9 | 60483 | 498 | GSE85066 |
| MCA20[6] | Mouse Cell Atlas (MCA) dataset of Microwell-seq data from >28 tissues (2-4 replicates each) and cultures | retaining only features that are shared across all replicates (of a given tissue), and observations for which metadata was available | 1-4 | 170 | >10,000 | >1,200,00 | figshare |
| Mereu20[7] | PBMC data from 13 platforms (Chromium, Chromium(sn), inDrop, C1HT-small and -medium, CEL-Seq2, ddSEQ, Drop-Seq, ICELL8, MARS-Seq, Quartz-Seq2, mcSCRB-Seq, and Smart-Seq2) | – | 13 | 9 | 23381 | 20237 | GSE133549 |
| Oetjen18[8] | Droplet-based scRNA-seq of bone marrow mononuclear cells from 20 healthy donors of different sex and age (25 samples in total) | removal of replicated samples (Ck, C1, C2, Sk1, Sk2, S1, S2) | 18 | – | 33694 | 72241 | GSE120221 |
| panc8 | eight human pancreatic islet cell datasets from five technologies (CEL-Seq, CEL-Seq2, inDrop (four replicates), Fluidigm C1, SMART-Seq2) | retaining the inDrop (technical) replicate with the highest number of cells | 5 | 13 | 23600 | 10963 | GSE81076, GSE85241, GSE86469, E-MTAB-5061 |
| TabulaMuris[9] | droplet-based scRNA-seq data from Mus musculus (8 male and female mice) across 20 organs and tissues | – | 10 | 13 | 23341 | 17404 | figshare |
| Tung17[10] | triplicated Fluidigm's C1 data of induced pluripotent stem cell (iPSC) lines of three individuals (9 samples in total) | – | – | 3 | 20327 | 864 | GSE77288 |
| Zheng17[11] | droplet-based scRNA-seq data of PBMCs from a single healthy individual | T cell subpopulations merged into CD4+ and CD8+ | – | 9 | 32738 | 68579 | SRP073767 |

**Supplementary Table 1:** Overview of reference datasets. Each entry specifies the dataset identifier, a brief description of the measurement technology and cell and/or tissue type, how data were filtered or preprocessed (if applicable), the number of batches (biological or technical replicates), clusters, features (genes or transcripts) and observations (cells), and the data source (E-X = ArrayExpress, GSE = Gene Expression Omnibus, SCP = Broad Institute's Single Cell Portal).

| **Dataset** | **Subset(s)** | **Type** | **Batch(es)** | **Cluster(s)** |
|---|---|---|---|---|
| CellBench | ✗ | b,k | 3 | 3 |
| | H2228 | b | 3 | H2228 |
| | celseq | k | sc_celseq | 3 |
| Ding20 | ✗ | b,k | 4 | 8 |
| | 10x.InhibNeuron | n | 10x Chromium | Inhibitory neuron |
| | ExcitNeuron | b | 4 | Excitatory neuron |
| | DroNcSeq | k | DroNc-seq | 5 |
| Gierahn17 | ✓ | n | 0 | 0 |
| Kang18 | ✗ | b,k | 8 | 8 |
| | 1015 | k | 1015 | 6 |
| | B | n | 1015 | B cells |
| | NK | n | 1015 | NK cells |
| Koh16 | ✓ | k | 0 | 7 |
| MCA20 | ✗ | b,k | 13 | 9 |
| | gland.AT2 | b | 4 | T cell_Cd8b1 high |
| | lung.AT2 | b | 4 | AT2 Cell |
| Mereu20 | ✗ | b,k | 13 | 9 |
| | CD4T | b | 13 | CD4 T cells |
| | ddSeq | k | ddSeq | 9 |
| Oetjen18 | ✓ | b | 18 | 0 |
| | R | n | R | 0 |
| panc8 | ✗ | b,k | 5 | 9 |
| | inDrop1.beta | n | indrop1 | beta |
| | inDrop.ductal | b | indrop1-4 | ductal |
| | SmartSeq2 | k | smartseq2 | 7 |
| TabulaMuris | ✗ | b,k | 10 | 31 |
| | limb.MSCs | n | Limb_Muscle | mesenchymal stem cell |
| | spleen | k | Spleen | 4 |
| Tung17 | ✓ | b | 3 | 0 |
| | NA19101 | n | NA19101 | 0 |
| Zheng17 | ✓ | k | 0 | 7 |
| | HSCs | n | 0 | HSCs CD34+ |
| | Monocytes | n | 0 | Monocytes CD14+ |

**Supplementary Table 2:** Overview of data subsets drawn to serve as references for simulation. Each entry specifies the dataset and subset identifier, type ($n$, $b$ or $k$), and the number or identity of the retained batch(es) and cluster(s) after filtering. Header rows list the dataset's original number of batches and clusters, and ✓ / ✗ indicate whether the complete dataset was included as a reference. In total, there are 10, 8, and 8 subsets of type $n$, $b$, and $k$, respectively.

### 2 Quality control summaries

Let $\mathbf{X}(\mathbf{Y}/\mathbf{Z}\ )_{G\times C}$ denote the count (expression) matrix of a reference or simulated dataset with genes $\mathcal{G}=\{g_1,\dots,g_G\}$ and cells $\mathcal{C}=\{c_1,\dots,c_C\}$ from batches $b=1,...,B$ and clusters $k=1,...,K$. Here, $B$ is the number of batches, $K$ is the number of clusters, $\mathbf{Y}$ and $\mathbf{Z}$ correspond to log1p-transformed counts per million (CPM) and log-library size normalized counts obtained with *scater*'s[12] *calculateCPM* and *logNormCounts*, respectively. For principal component-(PC-)based summaries, we ran *scran*'s[13] *modelGeneVar* (on $\mathbf{Z}$) and *getTopHVGs* to select the $n=500$ most highly variable features, and *scater*'s *calculatePCA* to compute their first $ncomponents=50$ PCs. The same set of inputs specified below were used to compute the respective summaries for all reference and simulated datasets.

| **Summary** | **Description/Interpretation** | **Formula/Implementation** |
|---|---|---|
| mean of logCPM | expression mean | $\mu=\frac{1}{C}\sum_{c=1}^{C}\mathbf{Y}_{gc}$ |
| variance of logCPM | expression variance | $\sigma=\frac{1}{C-1}\sum_{c=1}^{C}(\mathbf{Y}_{gc}-\mu)^2$ |
| coefficient of variation | expression variability relative its mean | $\sqrt{\sigma}/\mu$ |
| gene detection frequency | fraction of cells with non-zero count (for a given gene) | $\frac{1}{C}\sum_{c=1}^{C}\mathbb{1}(\mathbf{X}_{gc}\neq 0)$ |
| gene-to-gene-correlation | expression association between pairs of genes | $\frac{\text{cov}(\mathbf{Y}_g,\mathbf{Y}_{g'})}{\sigma_g\sigma_{g'}}$ |
| log-library size | log1p-transformed total counts | $\log(1+\sum_{g=1}^{G}\mathbf{X}_{gc})$ |
| cell detection frequency | fraction of detected genes (for a given cell) | $\frac{1}{G}\sum_{g=1}^{G}\mathbb{1}(\mathbf{X}_{gc}\neq 0)$ |
| cell-to-cell-correlation | expression association between pairs of cells | $\text{cov}(\mathbf{Y}_c,\mathbf{Y}_{c'})/(\sigma_c\cdot\sigma_{c'})$ |
| local density factor | relative measure of a cell's local density compared to those within its neighbourhood (in PCA space) | custom wrapper of functions from the *CellMixS* package[14] with PCs of $\mathbf{Z}$ as input |
| cell-to-cell distance | expression (dis)similarity between pairs of cells | Euclidean distance in PCA space of $\mathbf{Z}$ |
| KNN occurences | number of times a cell is a k-nearest neighbor (KNN) | *RANN*'s *nn2* function on PCs of $\mathbf{Z}$ with k set to 5% of cells |
| percent variance explained | fraction of expression variance accounted for by batch/cluster | *variancePartition*'s *fitExtractVarPartModel* function[15] with $\mathbf{Z}$ as input |
| silhouette width | similarity of a cell to its own group (batch/cluster) compared to others[16] | *cluster*'s *silhouette* function[17] on Euclidean distances in PCA space of $\mathbf{Z}$ |
| cell-specific mixing score | probability of being in an equally 'mixed' (same batch/cluster) neighborhood (in PCA space) | *CellMixS*'s *cms* function[14] with PCs of $\mathbf{Z}$ as input |

**Supplementary Table 3:** Overview of scRNA-seq data summaries used to compare reference and simulated data. Summaries are grouped by type: gene-, cell-level and global. For each summary, a brief description or possible interpretation is provided, as well as how it is computed (theoretically) or implemented (in R).

### 3 Method parameters

| | Estimation | Simulation |
|---|---|---|
| BASiCS | BASiCS_MCMC with MCMC sampler parameters N = 4000 iterations, thinning period Thin = 10, and burn-in period Burn = 2000; joint prior formulation for mean and over-dispersion (Regression = TRUE); and, using batches to estimate technical variability (WithSpikes = FALSE). | BASiCS_Sim with Mu_spikes = Phi = NULL, and other parameters passed from estimation. |
| ESCO | escoEstimate with default parameters, using raw counts as input, and cell group labels set to batch/cluster identifiers for type $b/k$. | escoSimulate with parameters passed from estimation, type = "single" for type $n$, and "group" otherwise. |
| hierarchicell | filter_counts with cells randomly split into two groups for type $n$ and $gene_thresh = cell_thresh = 0$ (i.e., retaining all genes and cells); compute_data_summaries with Raw = "raw" using raw counts as input; number of non-control groups and cells n_cases = cells_per_case = 1 (required, but removed prior to simulation); $n_controls$ and cells_per_control set to the number of reference batches and reference cells per batch, respectively; ncells_variation_type = "Fixed" (i.e., fixed number of cells per individual). | simulate_hierarchicell with parameters passed from estimation, and filtering for cells with Status == "Control" (i.e., removing non-control cells). |
| muscat | prepSim with min_size = NULL (i.e., retaining all subpopulation-sample combinations), and otherwise default parameters. | simData with dd = FALSE (i.e., no differentially distributed genes), and otherwise default parameters. |
| POWSC | Est2Phase with default parameters, using raw counts as input; called once for references of type $n$, and separately on each cluster for type $k$. | For type $n$, Simulate2SCE with perDE = 0 (i.e., no differentially expressed genes), and parameter estimates passed to both estParas1 and estParas2 (i.e., equivalent parameters for either group); for type $b/k$, SimulateMultiSCEs with multiProb set to the number of cells per batch/cluster in the reference, and parameter estimates passed to estParas_set. |
| powsimR | estimateParam with raw counts as input; RNAseq = "singlecell", Protocol = "UMI", Distribution = "NB", Normalisation = "scran", GeneFilter = 0 and SampleFilter = Inf (i.e., retaining all genes and cells for normalization and parameter estimation); number of group 1 cells n1 set to the number of reference cells, and of group 2 cells n2 = 2 (required, but removed prior to simulation); pDE = pLFC = 0 (no differential expression); nsims = 1 (one simulation replicate). | simulateDE with Normalisation = "scran" and DEmethod = "DESeq2", Counts = TRUE (i.e., input data corresponds to raw counts), and removing group 2 cells from the simulated count matrix. |
| scDD | preprocess with scran_norm = TRUE and condition set to a mock variable to randomly split reference cells into two groups; number differentially distributed genes nDE = nDP = nDM = nDB = nEP = 0 and nEE = 1 (i.e., only equivalently expressed genes); and, numSamples set to half the number of reference cells (since two groups are simulated). | simulateSet with parameters passed from estimation. |
| scDesign | – | design_data with total number of RNA-seq reads S set to the overall sum of reference counts, ncell set to the number reference cells, and ngroup = 1 (i.e., one cell state only). |
| scDesign2 | fit_model_scDesign2 with default parameters, using raw counts as input; cell identifiers set to cluster assignments for type $k$ and a (unique) mock identifier for type $n$. | simulate_count_scDesign2 with n_cell_new set to the number of reference cells, and cell_type_prop set to the frequency of each reference cluster. |
| SCRIP | For type $n$, splatEstimate with default parameters, using raw counts as input; no additional estimation for type $b$ and $k$. | For type $n$, SCRIPsimu with parameters passed from estimation; for type $b/k$, simu_cluster with CTlist set to unique reference batch/cluster identifiers; for all types, mode = "GP-trendedBCV". |
| SPARSim | SPARSim_estimate_parameter_from_data with default parameters, using both raw (raw_counts) and library size normalized counts (norm_data) as input; conditions set to batch identifiers for type $b$, and a mock variable (unique identifier for each cell) for type $n$. | SPARSim_simulation with parameters passed from estimation and output_batch_matrix = TRUE, extracting the result list's element count_matrix as output. |
| splatter | splatEstimate with default parameters. | splatSimulate with parameters passed from estimation. |
| SPSimSeq | – | SPsimSeq with n.genes/tot.samples set to the number of reference genes/cells, and model.zero.prob = genewiseCor = TRUE; for type $n$; batch set to a mock variable (1 for every cell) and batch.config = 1; for type $b$, additional filtering for genes with a total count of at least 10 in all batches (to prevent parameter estimation failure), batch set to reference cell batch assignments, and batch.config set to the frequency of each reference batch. |
| SymSim | BestMatchParams with raw counts as input, tech = "UMI", n_optimal = 1, and ngenes, ncells_total, nbatch set to the number of genes, cells, batches, in the reference. | SimulateTrueCounts with parameters passed from simulation and randseed = 1234; True2ObservedCounts with true_counts and meta_cell passed from SimulateTrueCounts's output, and gene_len sampled from the package's internal gene_len_pool data (uniformly and with replacement); for type $n/k$, DivideBatches with nbatch set to the number of reference batches/clusters and observed_counts_res set to True2ObservedCounts's output. |
| ZINB-WaVE | zinbFit with default parameters, and model matrix formula ~ batch/cluster for type $b/k$. | zinbSim with parameters passed from estimation. |
| zingeR | getDatasetZTNB with raw counts as input, reference cells randomly split into two groups, and pUp = 0 (i.e., no differentially expressed genes). | NBsimSingleCell with parameters passed from estimation. |

#### 4 Evaluation statistics

To evaluate how similar simulations are to the underlying (real) reference dataset, we compute both one- and two-dimensional tests on the similarity between quality control summaries (or pairs thereof) obtained from each reference-simulated dataset pair.

##### 4.1 One-dimensional

For every reference-simulation pair, we perform two tests on the similarity of summary distributions; these are briefly described below. Thus, we obtain one statistic per test, summary, method and reference dataset.

The **Kolmogorov–Smirnov (KS) test**[18] is a non-parametric test that may be used to quantify the distance between a pair of cumulative distribution functions (CDFs). Here, for each quality control summary, we compared the CDF obtained from reference ($F$) against simulated data ($G$). The KS statistic is defined as the largest absolute distance between $F$ and $G$, i.e.:

$$D(F,G) = sup_x|F(x) - G(x)|$$

Similar to the KS test, the **Wasserstein distance (w)** describes the distance between two distributions, where smaller/larger values correspond to more/less similarity; its exact value, however, is generally not straight-forward to interpret. The 1st Wasserstein metric is defined as:

$$W(F,G) = \left( \int_0^1 \left| F^{-1}(u) - G^{-1}(u) \right|^2 du \right)^{\frac{1}{2}}$$

where $F$ and $G$ denote the cumulative distribution function (CDF) of the reference and simulated dataset, respectively, and $F^{-1}$ and $G^{-1}$ their corresponding quantile functions.

KS tests were performed using the ks.test function with alternative = ‘two.sided‘ (i.e. under the null hypothesis, $F$ and $G$ are equal). In general, a KS statistic of $\approx 0$ suggests that reference and simulation are very similar; hence, smaller values correspond to better method performance.
To compute $W$, we used the wasserstein_metric function of the waddR package[19], which provides a (faster) Rcpp re-implementation of the original wasserstein1d function from the transport package[20].

##### 4.2 Two-dimensional

The **two-dimensional KS test** as proposed by Peacock[21] was performed using the peacock2 function of the Peacock.test package.

To compute the **Earth mover's distance (EMD)**[22] between a pair of quality control summaries $p$ and $q$ obtained from reference $x$ and simulated data $y$, we estimated a two-dimensional kernel density over the range of observed values, i.e. $[\min(p_x, p_y), \max(p_x, p_y)]$ and $[\min(q_x, q_y), \max(q_x, q_y)]$, using MASS' kde2d function[23] with $n = 25$ grid points. The EMD was then computed using emdist's emd2d function[24], and divided by $n$ to make results independent of the number of evaluations points.

Both, the two-dimensional KS test and EMD were computed for each relevant pair of summaries, resulting in nine statistics per test, method, and reference dataset. Cell-to-cell distance and correlation, as well as gene-to-gene correlation were excluded from two-dimensional evaluations because they were computed for a random subset of gene- and cell-pairs, respectively. Similarly, the PVE, LDF, CMS and silhouette width were not included because they aim to capture global structure, and are expected to be unrelated to other metrics. Thus, we consider the following pairs of summaries:

| | variance of logCPM | gene detection frequency | coefficient of variation | cell detection frequency | KNN occurrences |
|---|---|---|---|---|---|
| average of logCPM | ✓ | ✓ | ✓ | | |
| variance of logCPM | | ✓ | ✓ | | |
| gene detection frequency | | | ✓ | | |
| log-library size | | | | ✓ | ✓ |
| cell detection frequency | | | | | ✓ |

### 5 Downstream

If not mentioned otherwise, all functions were run using default parameters. Throughout, $K$ corresponds to the 'true' number of clusters; logcounts correspond to log-transformed library size normalized counts obtained with scater's normalizeCounts function; and, principal component analysis and principal components are abbreviated with PCA and PCs, respectively.

#### 5.1 Integration

Integration methods were implemented as in Chazarra-Gil *et al.*[25], including ComBat[26], Harmony[27], fastMNN and mnnCorrect[28], limma[29], and Seurat[30]. To evaluate method performances, cell-specific mixing scores (CMS) and the difference in local density factors (ΔLDF) were computed using the cms and ldfDiff function, respectively, of the CellMixS package[14].

To make metrics comparable, we: i) subtracted 0.5 to center CMS at 0 (denoted CMS*); and, ii) centered (at 0) and scaled (to range 1) ΔLDF (denoted ΔLDF*). From these, we computed the Batch Correction Score (BCS) as the sum of |CMS*| and |ΔLDF*|. Thus, for all three metrics, a value of 0 indicates 'good' mixing for a given cell. When aggregating results (e.g., for heatmap visualizations), metrics were first averaged across cells within each batch and, secondly, across batches, in order to give equal weight to each batch independent to its size and complexity.

#### 5.2 Clustering

Clustering methods were implemented as in Duò *et al.*[31], including CIDR[32], hierarchical clustering (HC) and k-means[33] (KM) on PCA, pcaReduce[34], SC3[35], Seurat[30], TSCAN[36], and KM on t-SNE[37]. If applicable, the number of clusters was set to match the number of true (annotated respective simulated) clusters. To evaluate the performance of each method, we matched true and predicted cluster labels using the Hungarian algorithm[38], and computed cluster-level recall, precision, and F1 score (the harmonic mean of precision and recall):

$$\text{recall} = \frac{TP}{FP + FN}$$

$$\text{precision} = \frac{TP}{TP + FP}$$

$$\text{F1 score} = \frac{2 \cdot TP}{2 \cdot TP + FP + FN}$$

### 6 Computational workflow

For this benchmark, we designed a *Snakemake* workflow[39] that: i) reproducibly retrieves publicly available scRNA-seq datasets; ii) runs a set of simulation methods on subsets drawn from each reference dataset; iii) computes various global, gene- and cell-level quality control summaries; iv) compares summaries and relevant pairs thereof between reference and simulated data; v) quantifies parameter estimation and data simulation runtimes; vi) compares performance of methods for integration and clustering between reference and simulated data; and, vii) generates a variety of visualizations to consolidate results. The *Snakemake* is structured as follows:

- *config.yaml* specifies the R library and version to use
- *code/* contains all R scripts used in the workflow
- *data/* contains raw, subsetted, filtered and simulated scRNA-seq datasets, as well as simulation parameter estimates
- *meta/* contains two .json files that specify simulation method (*methods.json*) and reference subset (*subsets.json*) configurations
- *outs/* contains all results from computations (as .rds files), typically *data.frames*
- *plts/* contains all visual outputs (as .pdf files), and corresponding *ggplot* objects (as .rds files) for subsequent arrangement into 'super'-figures
- *figs/* contains figures (as *.pdf* files) that combine various content-related .rds objects from *plts/*

#### 6.1 Preprocessing

##### 6.1.1 Retrieval

We retrieved each reference dataset from a publicly accessible source, e.g., GitHub, Bioconductor's *ExperimentHub* or the Gene Expression Omnibus (GEO) database, using a self-contained and reproducible R script. Raw data are initially formatted into *SingleCellExperiment* objects[40], but left unprocessed otherwise. In rare cases (for example, *Kang18*), we apply some filtering to, e.g., remove unassigned cells, multiplets, and samples that have undergone experimental treatment (see Supp. Tab. 1).

##### 6.1.2 Filtering

As insufficient numbers of cells and low-quality observations can interfere with estimation of simulation parameters, we filter each reference dataset to: i) remove group-instances with fewer than 50 cells (here, groups correspond to batches or clusters, depending on the reference dataset's complexity); ii) retain genes with a count greater than 1 in at least 10 cells; and, iii) retain cells with at least 100 detected genes, i.e. non-zero counts.

##### 6.1.3 Subsetting

Finally, we draw various subsets from each reference dataset according to a configuration file (*meta/subsets.json*) that specifies, for each subset, which groups (i.e. batches or clusters) to retain, and (optionally) the number of genes and cells to downsample to. This gives rise to a set of subsets that serve as references for simulation (see Supp. Tab. 2).

The number of drawn subsets may vary from dataset to dataset, and is dependent on how many batches and/or clusters the reference provides. In general, we retain one subset per type for each reference, i.e., a type $n$, $b$ and/or $k$ subset (if there are multiple batches and/or clusters). For complex datasets (large number of genes/cells, batches/clusters), we preferably select for large and cleanly annotated groups.

#### 6.2 Simulation

Using a separate configuration file (*meta/methods.json*), we tag methods according to the features they can accommodate, i.e., one or many of batches ($b$), clusters ($k$), or neither ($n$). Each method is then run on all references that match the supported type(s). Thus, from each (real) reference dataset, we obtain corresponding simulated data for each method (see Tab. 1).

#### 6.3 Quality control

For each dataset, we compute the quality control summaries detailed in Supp. Tab. 3, giving rise to a set of global, gene- and cell-level summaries per (reference) dataset and (simulated) dataset-method pair.

#### 6.4 Performance evaluation

Next, we perform one- (and two-dimensional) tests on the similarity of reference-simulation summary pairs (and relevant pairs of summaries), resulting in a corresponding set of 14 and 9 statistics (per test) for one- and two-dimensional comparisons, respectively.

#### 6.5 Consolidation of results

##### 6.5.1 Filtering

All gene-level summaries as well as cell-level summaries that include pairwise computations (i.e., cell-to-cell distance and correlation) or capture global structure (i.e., local density factor and KNN occurrences) are dependent on the subset of cells they are computed on. Thus, we retain both group-level (per batch/cluster) as well as global (all cells) results for these summaries with one exception: gene-to-gene correlation was evaluated globally only (since we expect gene-to-gene correlation to be most interpretable across all cells in a dataset). For all other gene- and cell-level summaries, only group-level results are retained. Thus, we keep the following statistics for evaluation and visualization (○ = not computed):

| | average of logCPM | variance of logCPM | gene detection frequency | coefficient of variation | gene-to-gene correlation | log-library size | cell detection frequency | cell-to-cell correlation | local density factor | cell-to-cell distance | KNN occurrences | percent variance explained | cell-specific mixing score | silhouette width |
|---|---|---|---|---|---|---|---|---|---|---|---|---|---|---|
| global | ✓ | ✓ | ✓ | ✓ | ✓ | ✗ | ✗ | ✓ | ✓ | ✓ | ✓ | ✓ | ✓ | ✓ |
| group-level | ✓ | ✓ | ✓ | ✓ | ✗ | ✓ | ✓ | ✓ | ✓ | ✓ | ✓ | ○ | ○ | ○ |

##### 6.5.2 Averaging

When aggregating results (e.g., heatmaps), we first average statistics across global and/or group-level results (depending on the summary) with equal weights. Secondly, in order to weight datasets equally (independent of the number of drawn subsets), we first average statistics across subsets and, lastly, across datasets.

##### 6.5.3 Ranking

Methods/summaries are ranked according to their average statistic across summaries/methods, with equal weights given to all variables. For all statistics, lower values indicate better performance (i.e., 0 = best, 1 = worst).

### 7 Supplementary data

#### 7.1 QC summaries

obj-qc_ref/sim.rds are data.frames containing gene-, cell-level, and global quality control (QC) summaries across all references and methods. Specifically, these include:

- datset, subset: reference dataset and subset identifier
- metric: gene-, cell-level, or global quality control summary
- method: simulation method used ('ref' for the non-synthetic reference dataset)
- group: cell grouping used (one of 'global', 'batch', or 'cluster')
- id: cell group identifier, e.g., the batch or cluster annotation ('foo' when 'group' is 'global')
- value: summary value for a given feature (gene-level), cell (global and cell-level), or pair thereof (e.g., correlations)

#### 7.2 1/2D statistics

obj-stat_1/2d.rds are data.frames containing one-/two-dimensional test statistics results across all datasets, methods, and summaries (or relevant pairs thereof). In particular, these comprise:

- method: simulation method used to generate the data
- stat1/2d: test statistic used for comparing reference and simulation summary (or summaries) ('ks(2)' for (2D) Kolmogorov-Smirnov, 'ws' for Wasserstein metric, 'emd' for earth mover's distance)
- datset, subset: reference dataset and subset identifier
- metric(1,2): gene-, cell-level, or global quality control summary (or summaries)
- group: cell grouping used (one of 'global', 'batch', or 'cluster')
- id: cell group identifier, e.g., the batch or cluster annotation ('foo' when 'group' is 'global')
- stat: value of the test statistic

#### 7.3 Integration results

obj-batch_res.rds is a data.frame containing integration results for reference and simulated data across all type $b$ datasets and methods, and integration methods. Specifically, it includes the following columns:

- datset, subset: reference dataset and subset identifier
- method: simulation method used ('ref' for the non-synthetic reference dataset)
- batch_method: integration method used to correct for batch effects
- batch: ground-truth cell batch label
- ldf, cms: cell-specific difference in local density factor and mixing score

#### 7.4 Clustering results

obj-clust_res.rds is a data.frame containing clustering results for reference and simulated data across all type $k$ datasets and methods, and clustering methods. Specifically, it includes the following columns:

- datset, subset: reference dataset and subset identifier
- method: simulation method used ('ref' for the non-synthetic reference dataset)
- clust_method: clustering method used to predict cell cluster assignments
- cluster: ground-truth cell cluster annotation
- pr, re, F1: cluster-level precision, recall, and F1 score

#### 7.5 Runtimes

obj-rts.rds is a data.frame containing timings of parameter estimation and data simulation across all methods, and 5 replicates each for various random gene- and cell-subsets of one dataset per type. It includes:

- method: simulation method used
- datset, subset: reference dataset and subset identifier
- reftyp: reference dataset type (one of 'n', 'b', 'k', or 'g')
- ngs, ncs: number of genes/cells samples ('NA' if no downsampling)
- est, sim: runtime (in seconds) for parameters estimation and data simulation
  ('Inf' when estimation/simulation failed, 'est' is 'NA' when there is no separate estimation step)
