## Supplementary Figures for "Built on sand: the shaky foundations of simulating single-cell RNA sequencing data"

### 1 Dimension reduction

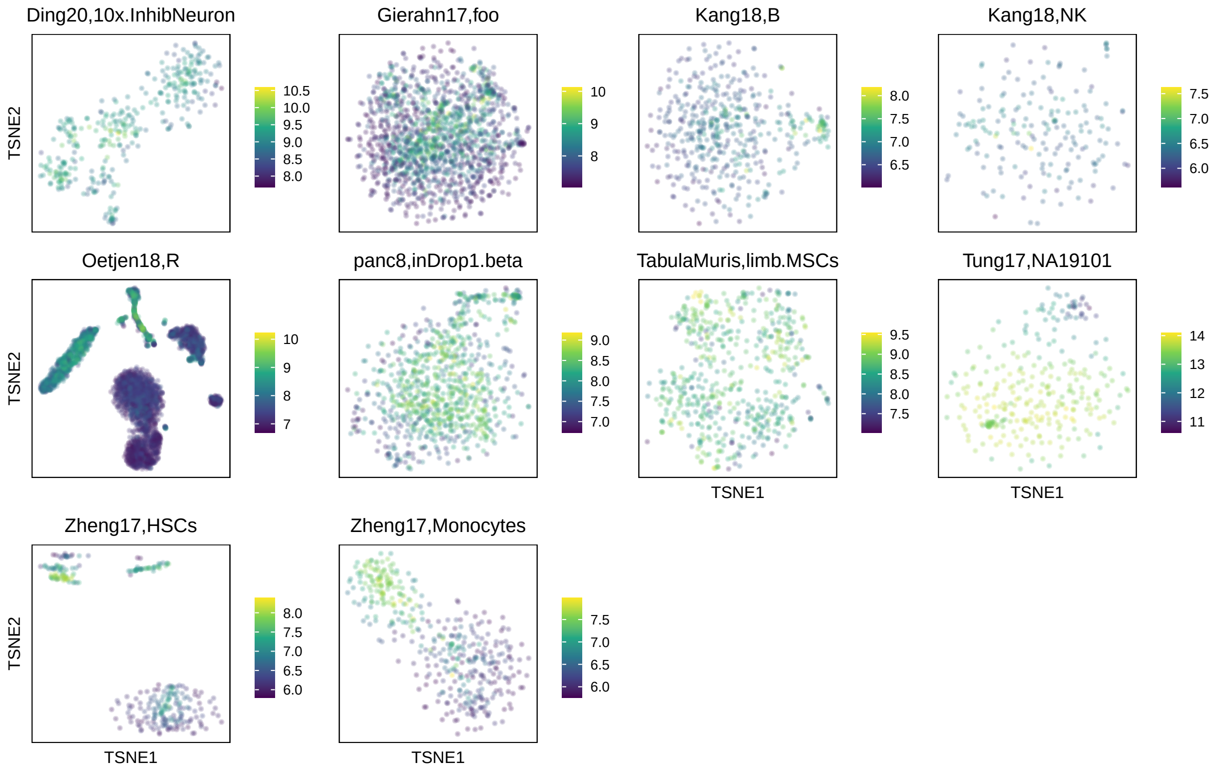

**Supplementary Figure 1:** t-SNE plots of type $n$ datasets. Cells (points) are colored by log1p-transformed library size (total counts). Plot titles correspond to comma-separated reference and subset identifiers.

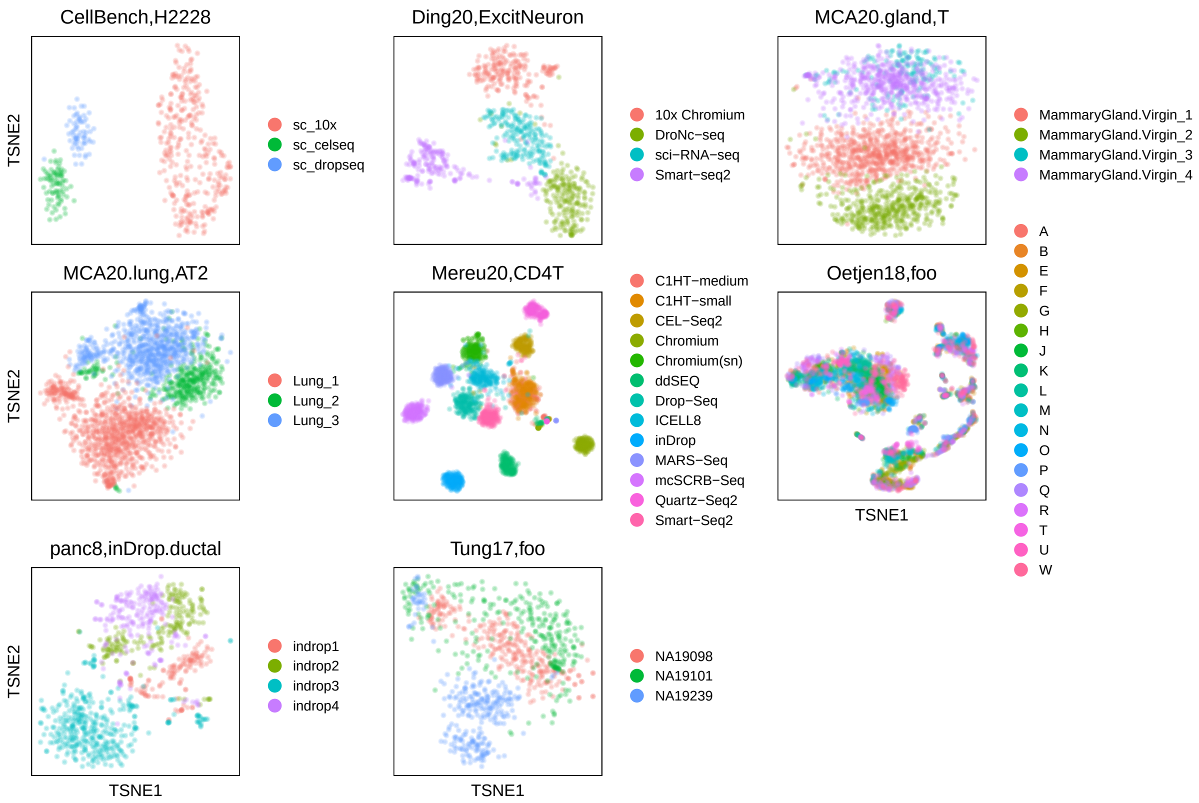

**Supplementary Figure 2:** t-SNE plots of type $b$ datasets. Cells (points) are colored by batch (biological or technical replicate). Plot titles correspond to comma-separated reference and subset identifiers.

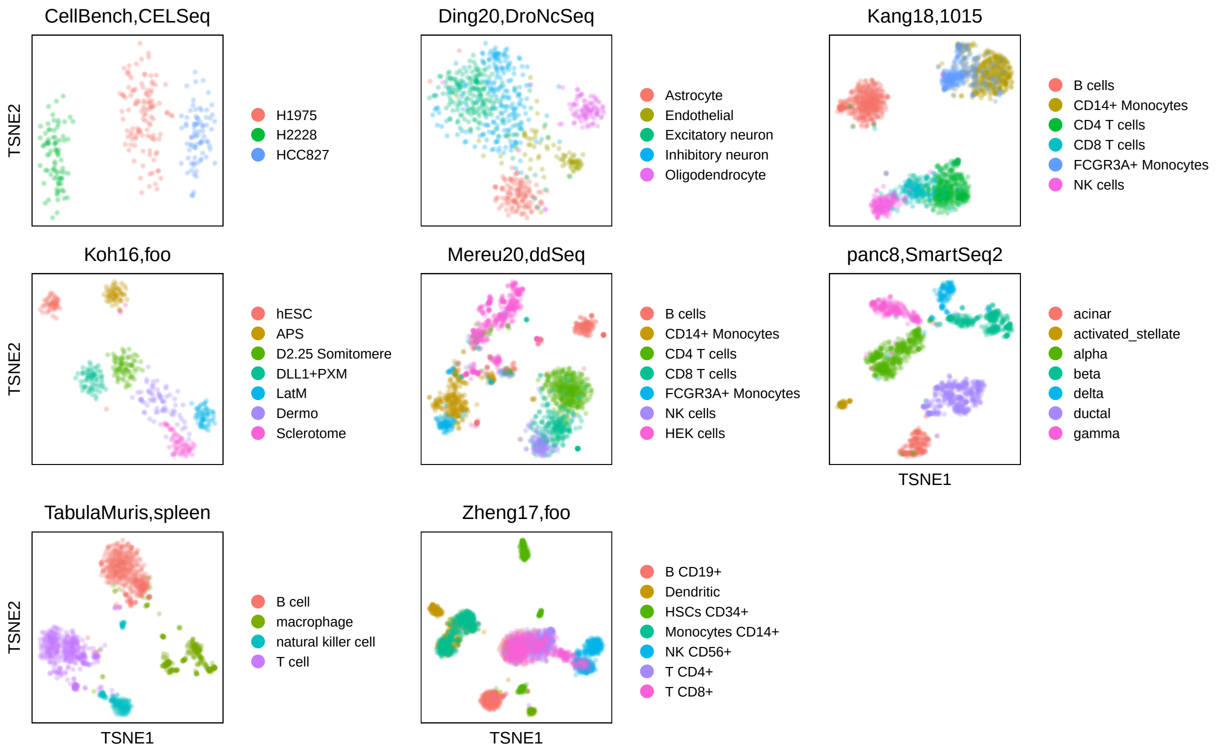

**Supplementary Figure 3:** t-SNE plots of type $k$ datasets. Cells (points) are colored by cluster (cell subpopulation). Plot titles correspond to comma-separated reference and subset identifiers.

### 2 Evaluation statistics

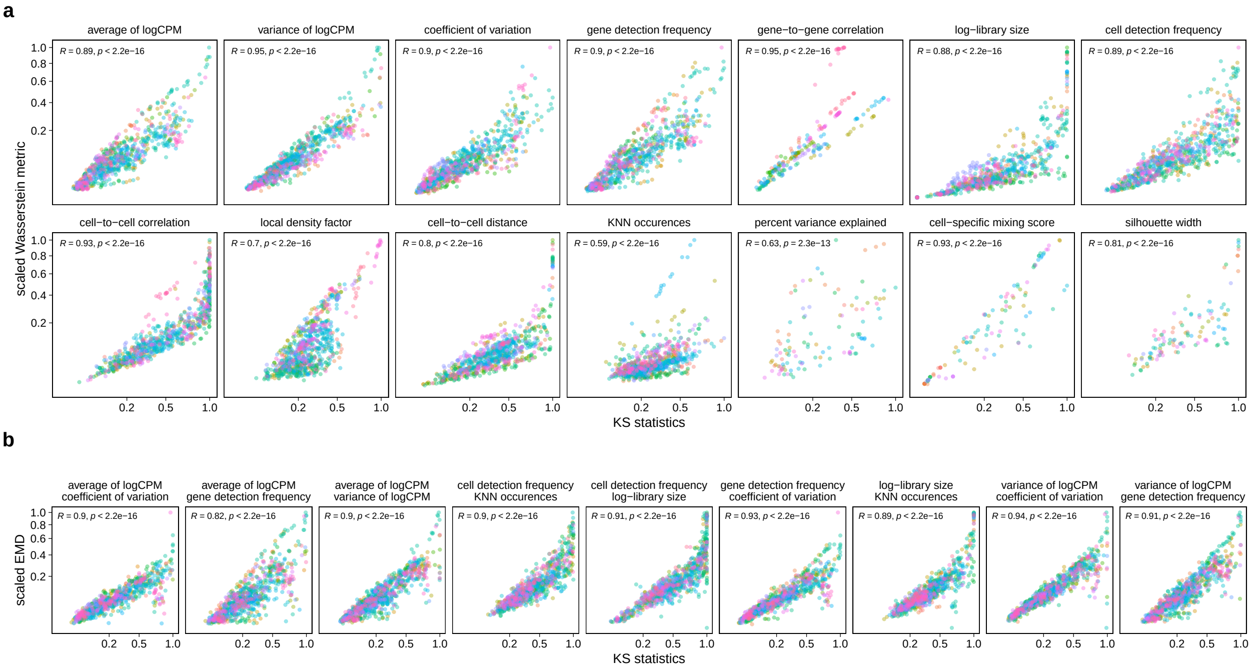

**Supplementary Figure 4:** Comparison of (a) one- and (b) two-dimensional test statistics. Each point corresponds to test statistics obtained for a given metric, method, dataset, and (if applicable) group (i.e., batch or cluster); points are colored by dataset. For comparability, Wasserstein metric and EMD values (y-axes) are scaled between 0 and 1 for each metric (panel). For datasets other than type $n$, only batch- and cluster-level (not global) results are included. Annotations represent Pearson correlation coefficients ($R$) and corresponding p-values ($p$).

#### 2.1 One-dimensional

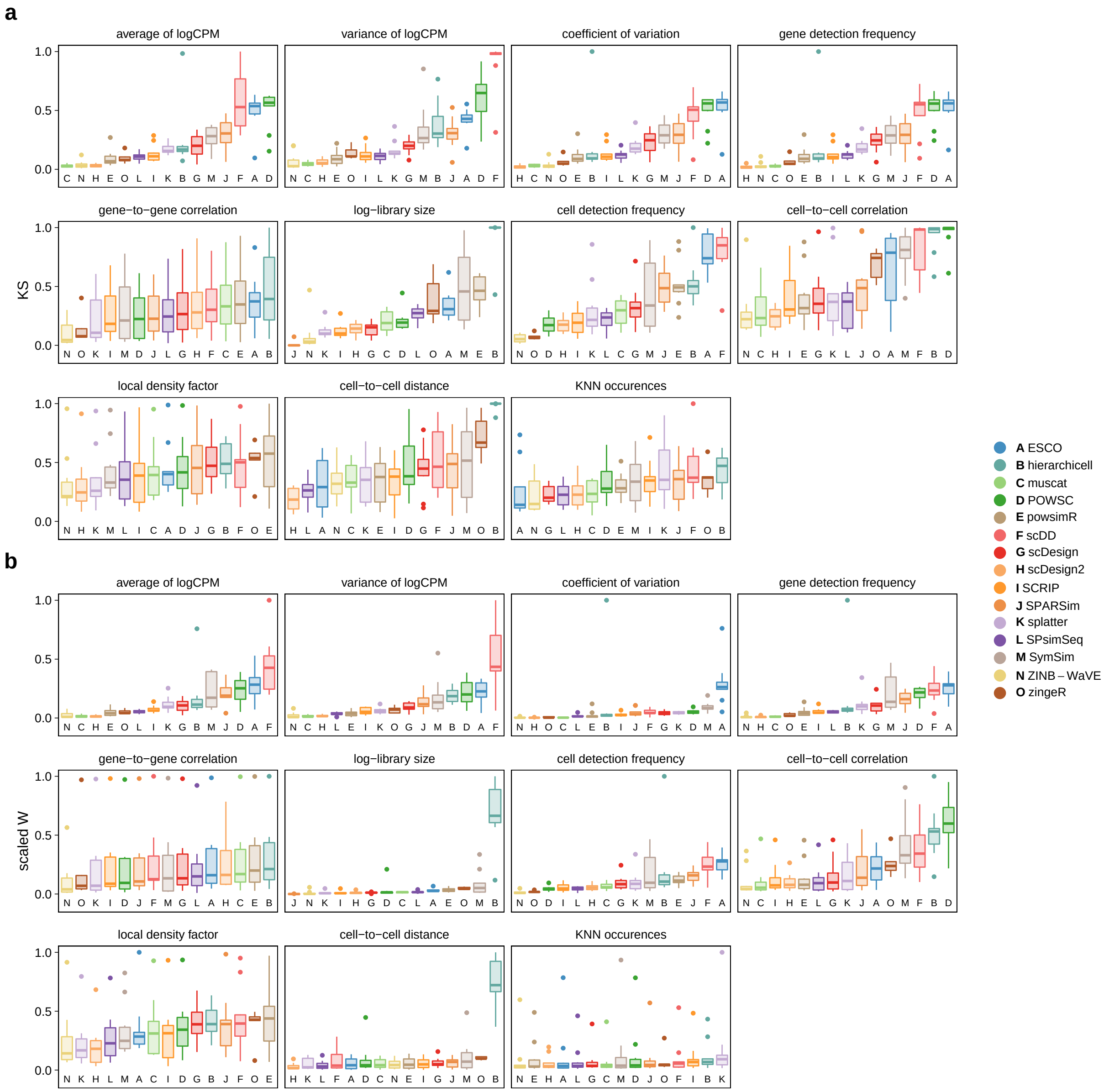

**Supplementary Figure 5:** Comparison of KS statistics (a) and Wasserstein metrics (b), type $n$. For each metric (panel), methods (x-axis) are ordered according to their average.

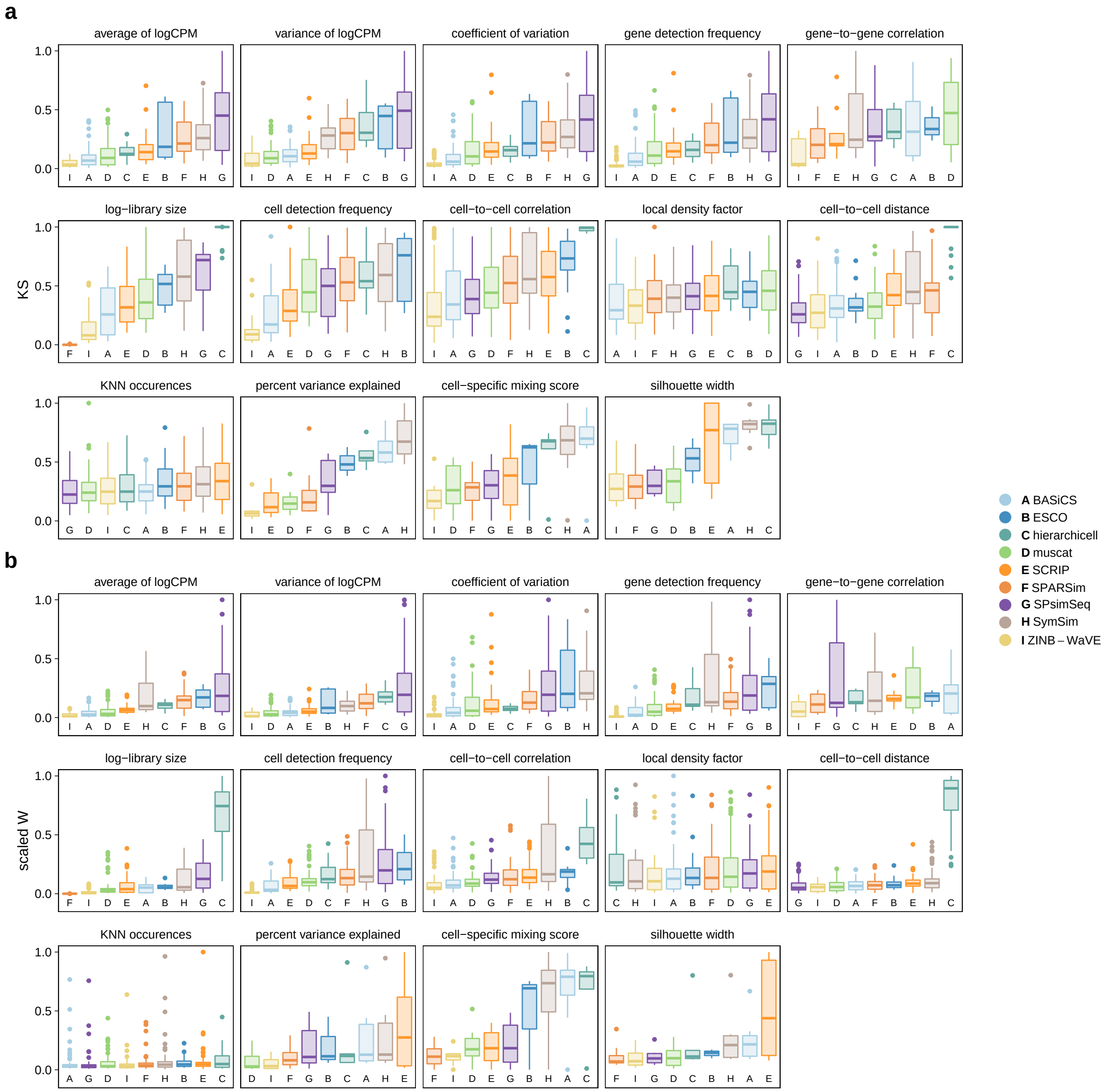

**Supplementary Figure 6:** Comparison of KS statistics (a) and Wasserstein metrics (b), type $n$. For each metric (panel), methods (x-axis) are ordered according to their average.

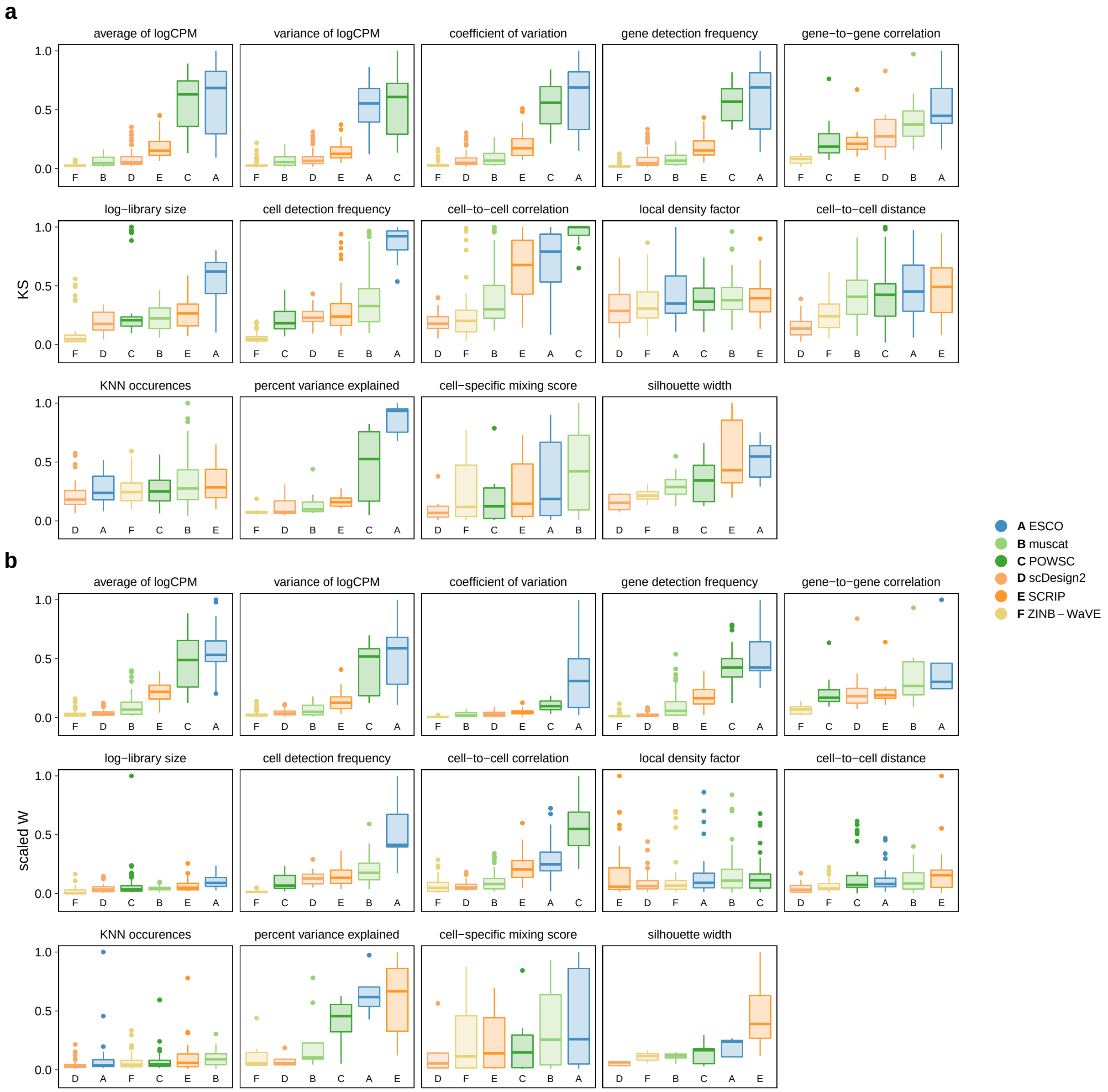

**Supplementary Figure 7:** Comparison of KS statistics (a) and Wasserstein metrics (b), type *b*. For each metric (panel), methods (x-axis) are ordered according to their average.

#### 2.2 Two-dimensional

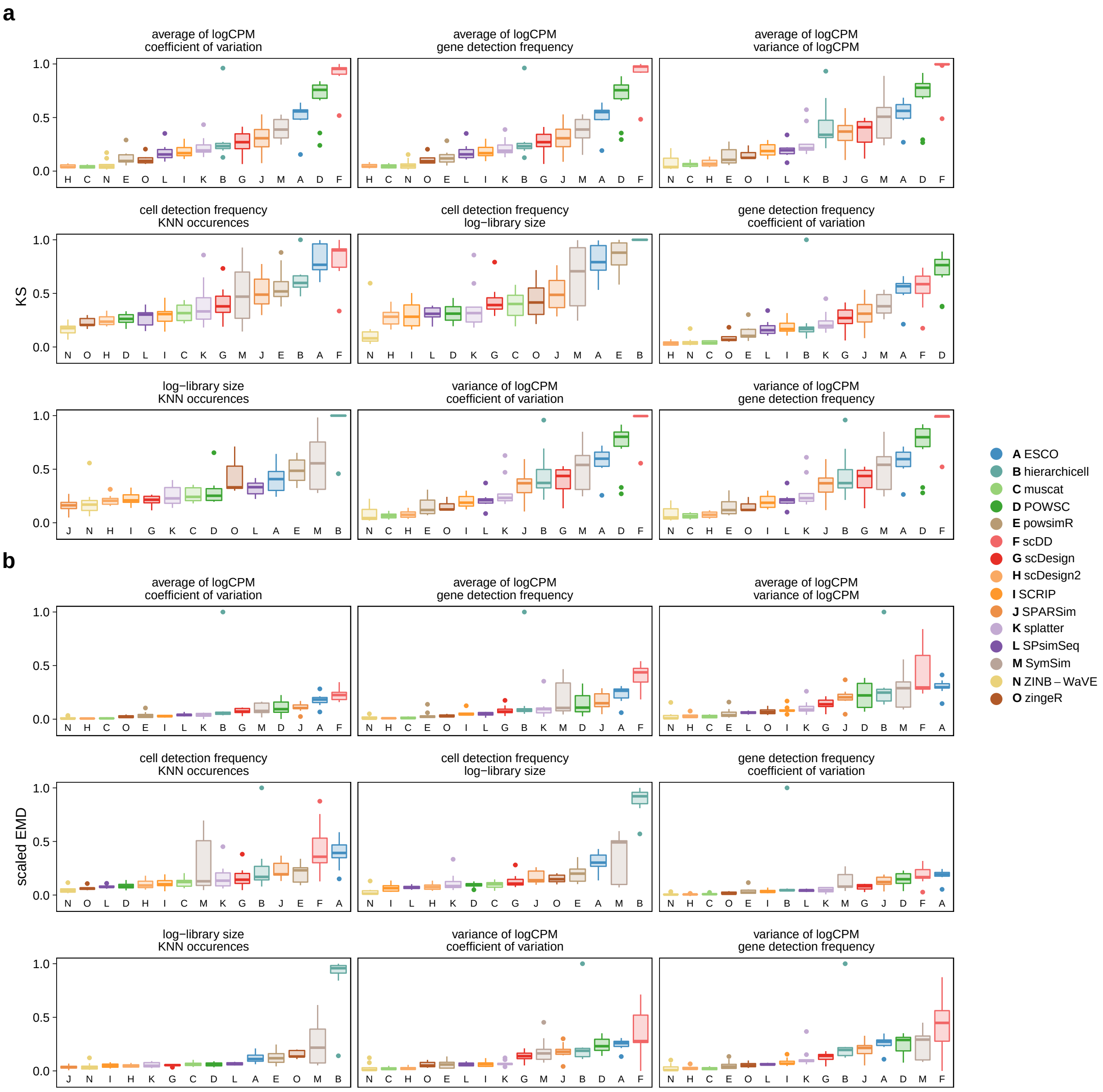

**Supplementary Figure 8:** Comparison of KS statistics (a) and EMDs (b), type $n$. For each pair of metrics (panel), methods (x-axis) are ordered according to their average.

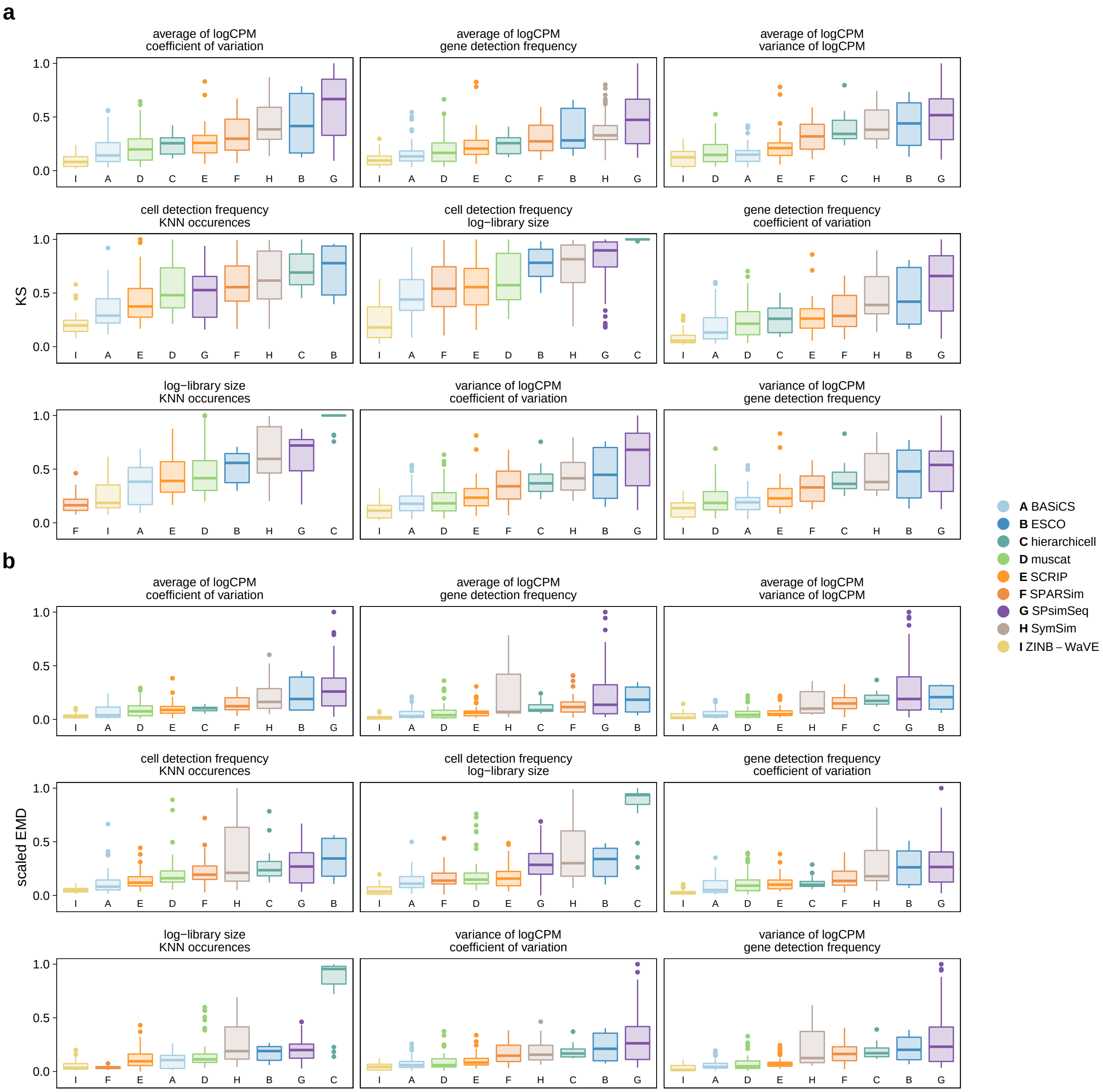

**Supplementary Figure 9:** Comparison of KS statistics (a) and EMDs (b), type *b*. For each pair of metrics (panel), methods (x-axis) are ordered according to their average.

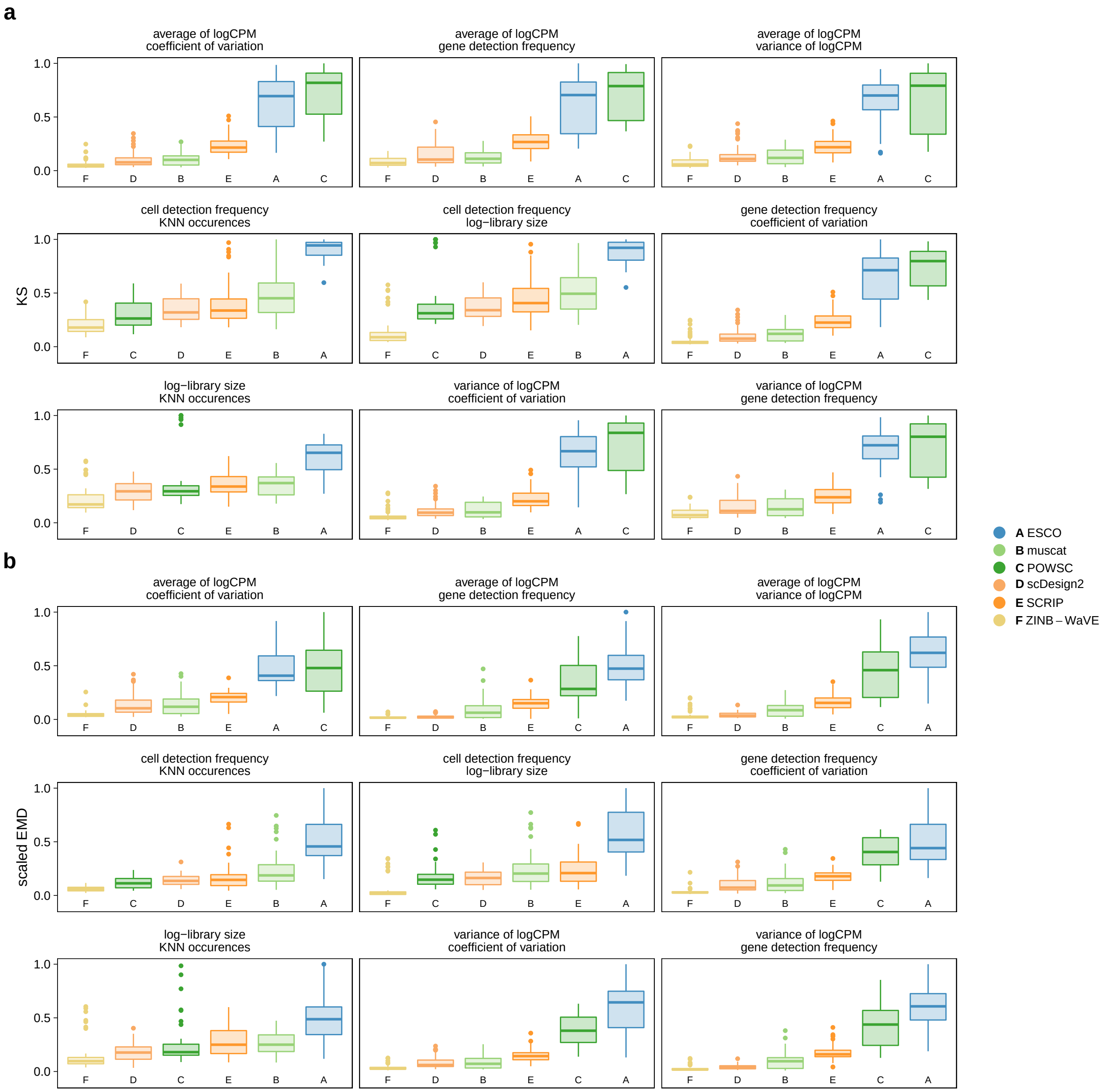

**Supplementary Figure 10:** Comparison of KS statistics (a) and EMDs (b), type $k$. For each pair of metrics (panel), methods (x-axis) are ordered according to their average.

### 3 Scalability

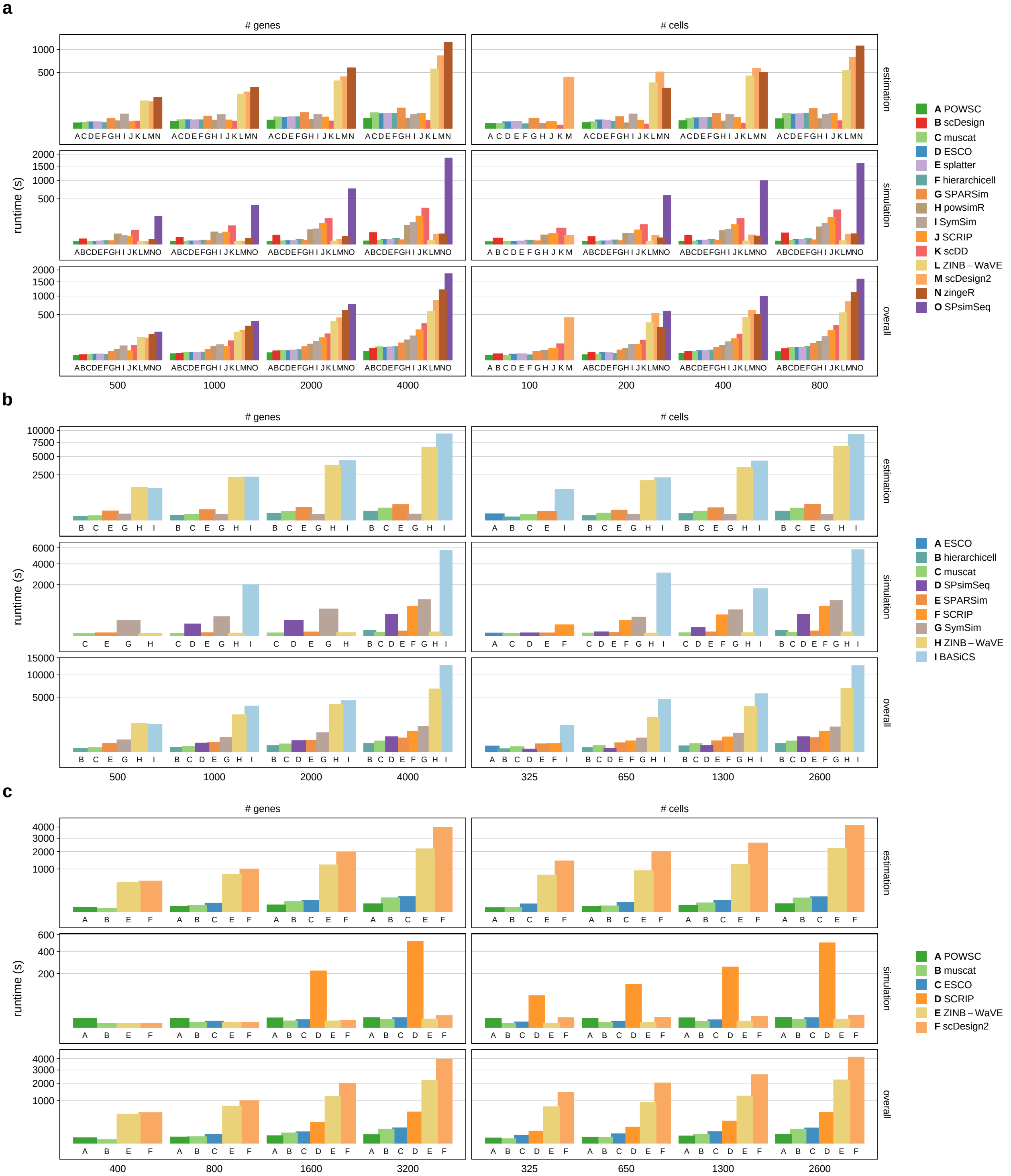

**Supplementary Figure 11:** Comparison of parameter estimation (optional), data simulation, and overall runtimes; stratified by type: (a) $n$, (b) $b$, and (c) $k$. Bars correspond to runtimes in seconds (s) averaged across 5 replicates per number of genes and cells, respectively. For each replicate, the number of genes (cells) was fixed when downsampling cells (genes). Methods are ordered by their average overall runtime across all gene and cell subsets.

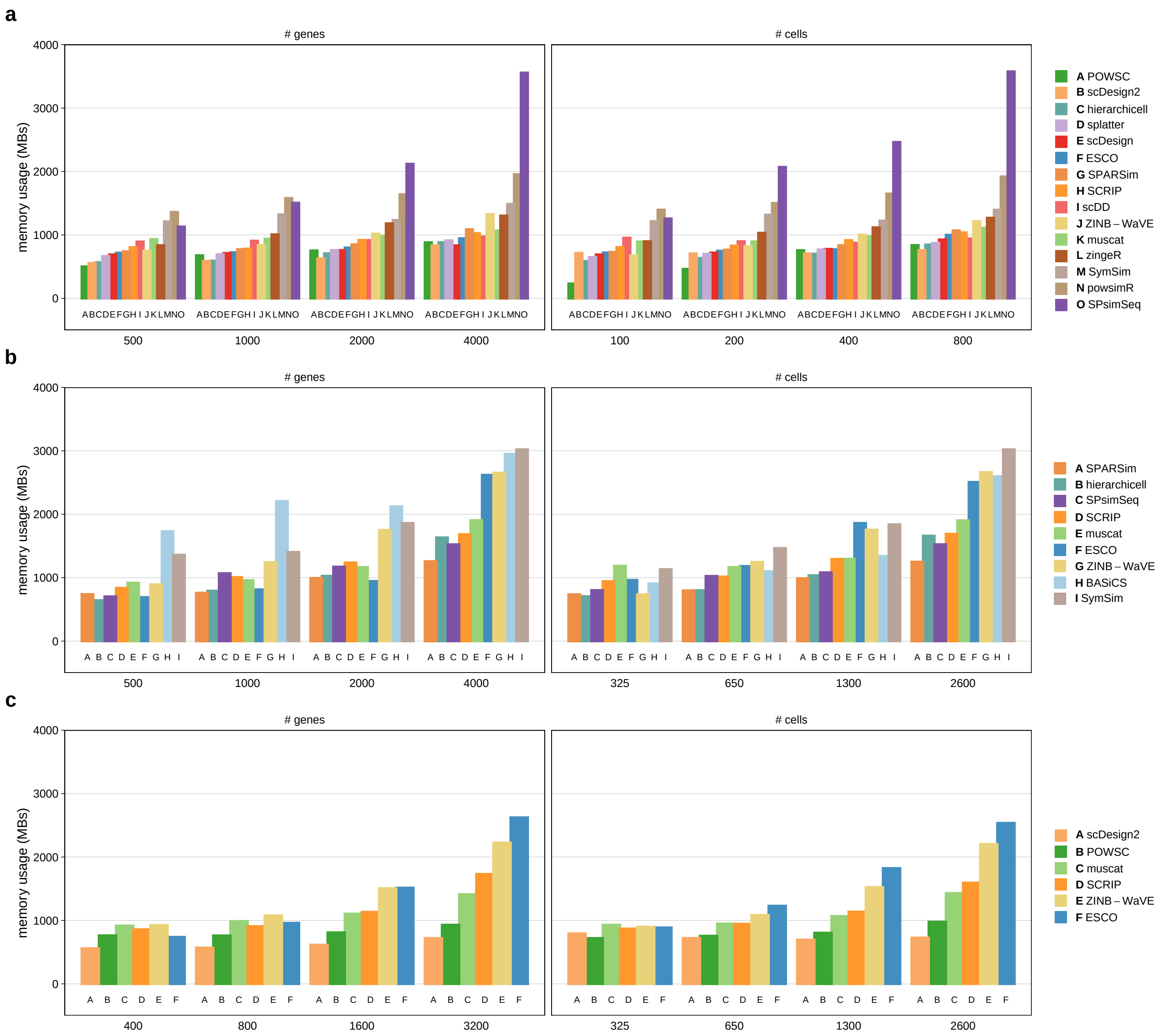

**Supplementary Figure 12:** Comparison of memory usage; stratified by type: (a) $n$, (b) $b$, and (c) $k$. Bars correspond to Resident Set Size (RSS) in megabytes (MBs) averaged across 5 replicates per number of genes and cells, respectively. For each replicate, the number of genes (cells) was fixed when downsampling cells (genes). Methods are ordered by their average memory usage across all gene and cell subsets.

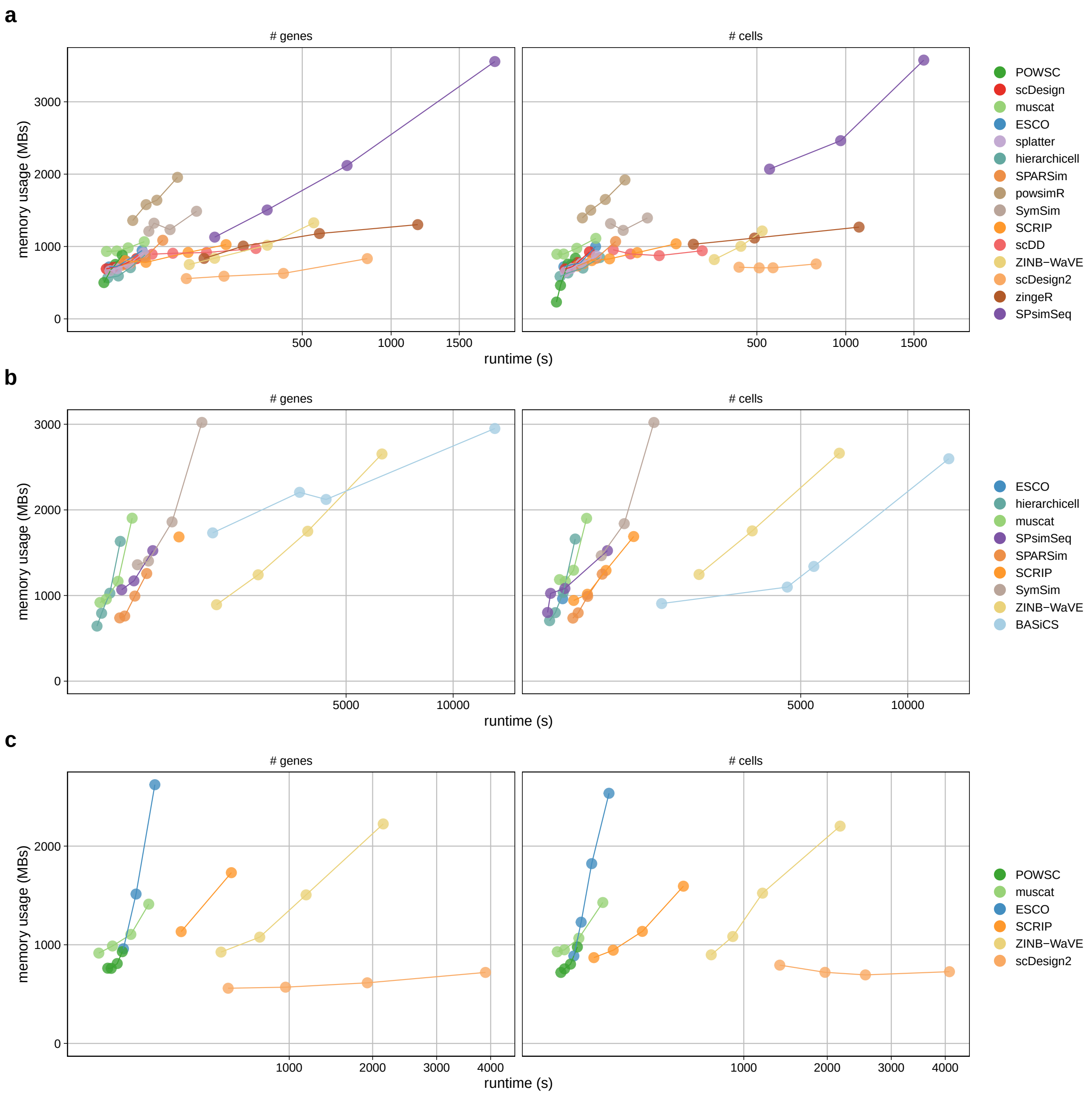

**Supplementary Figure 13:** Runtime vs. memory usage; stratified by type: (a) $n$, (b) $b$, and (c) $k$. Each point corresponds to overall (i.e., parameter estimation and simulation) runtime (s) and Resident Set Size (RSS) in megabytes (MBs) averaged across 5 replicates for a given gene and cell subset, respectively. For each replicate, the number of genes (cells) was fixed when downsampling cells (genes).

### 4 Downstream

#### 4.1 Integration

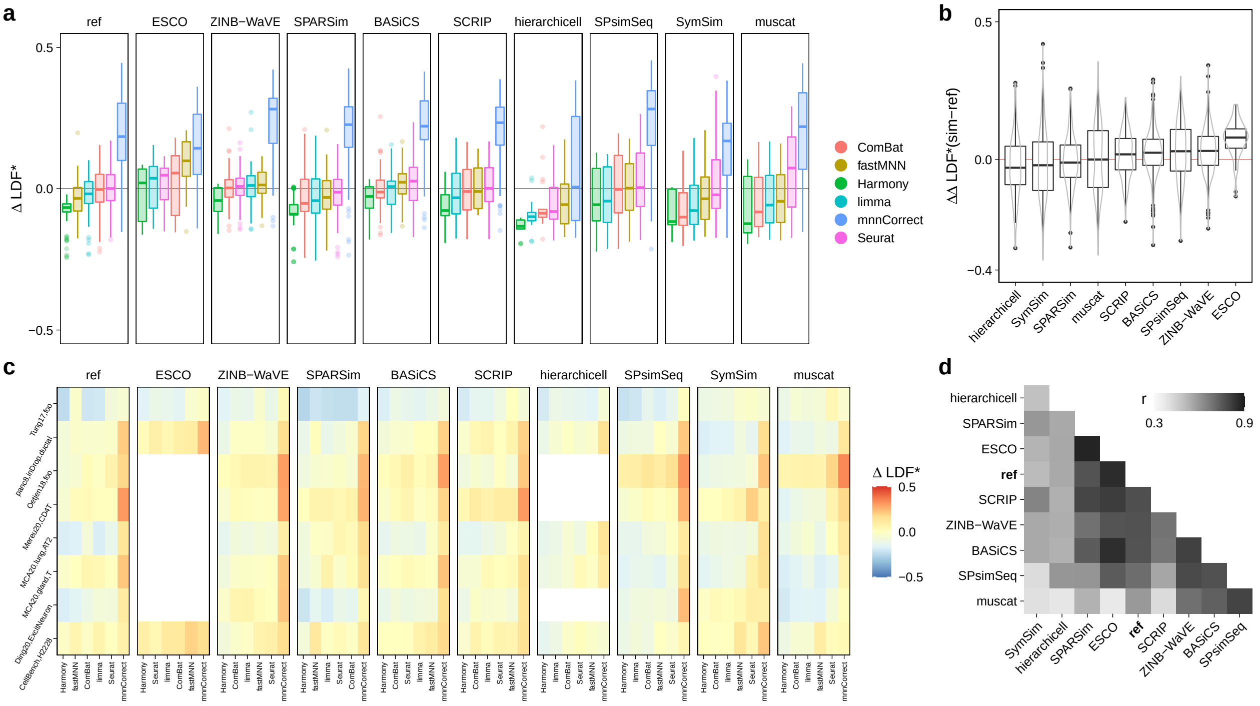

**Supplementary Figure 14:** Comparison of integration results across (experimental) reference and (synthetic) simulated data. (a) Boxplot of batch-level difference in local density factors ($\Delta$LDF) across all type $b$ references, simulation and integration methods. (b) Boxplot of difference ($\Delta$) in batch-level $\Delta$LDF obtained from *ref*erence and *sim*ulated data. (c) Heatmap of dataset-level $\Delta$LDF across integration methods (columns) and datasets (rows), stratified by simulator (panels). (d) Heatmap of Spearman's rank correlation ($\rho$) between $\Delta$LDF across datasets and integration methods.

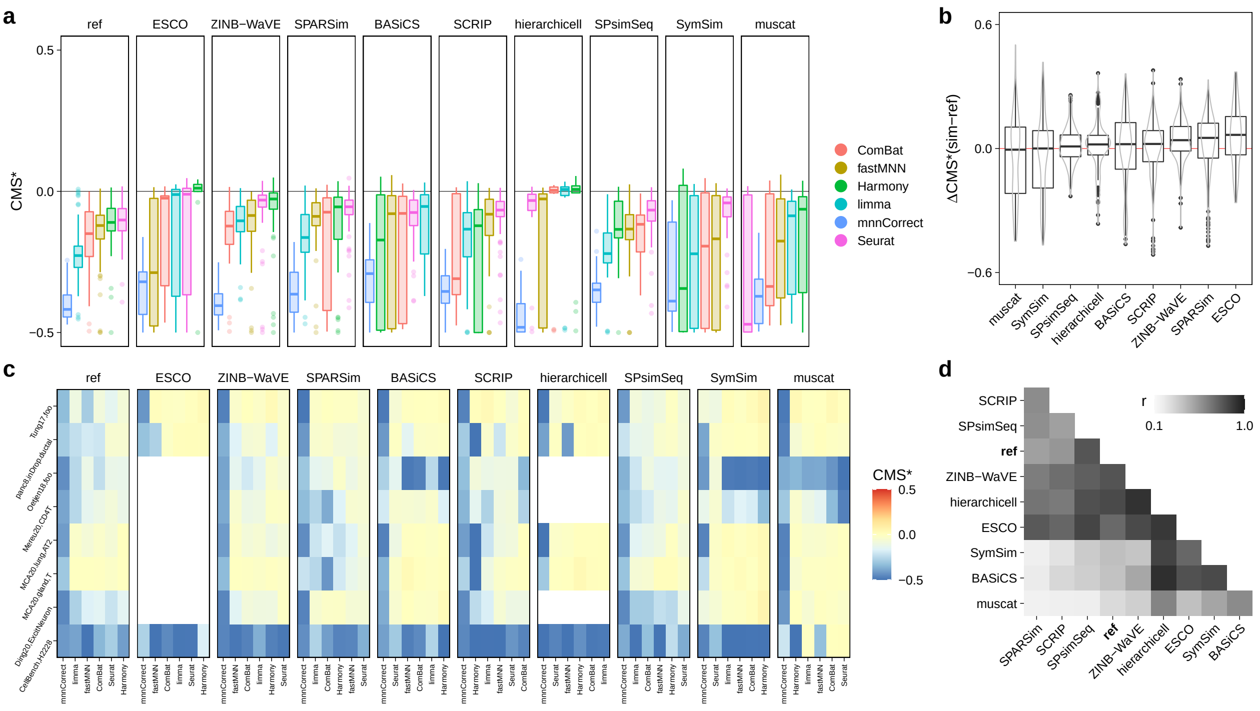

**Supplementary Figure 15:** Comparison of integration results across (experimental) reference and (synthetic) simulated data. (a) Boxplot of batch-level cell-specific mixing scores (CMS) across all type *b* references, simulation and integration methods. (b) Boxplot of difference (Δ) in batch-level CMS obtained from *ref*erence and *sim*ulated data. (c) Heatmap of dataset-level CMS across integration methods (columns) and datasets (rows), stratified by simulator (panels). (d) Heatmap of Spearman's rank correlation ($\rho$) between CMS across datasets and integration methods.

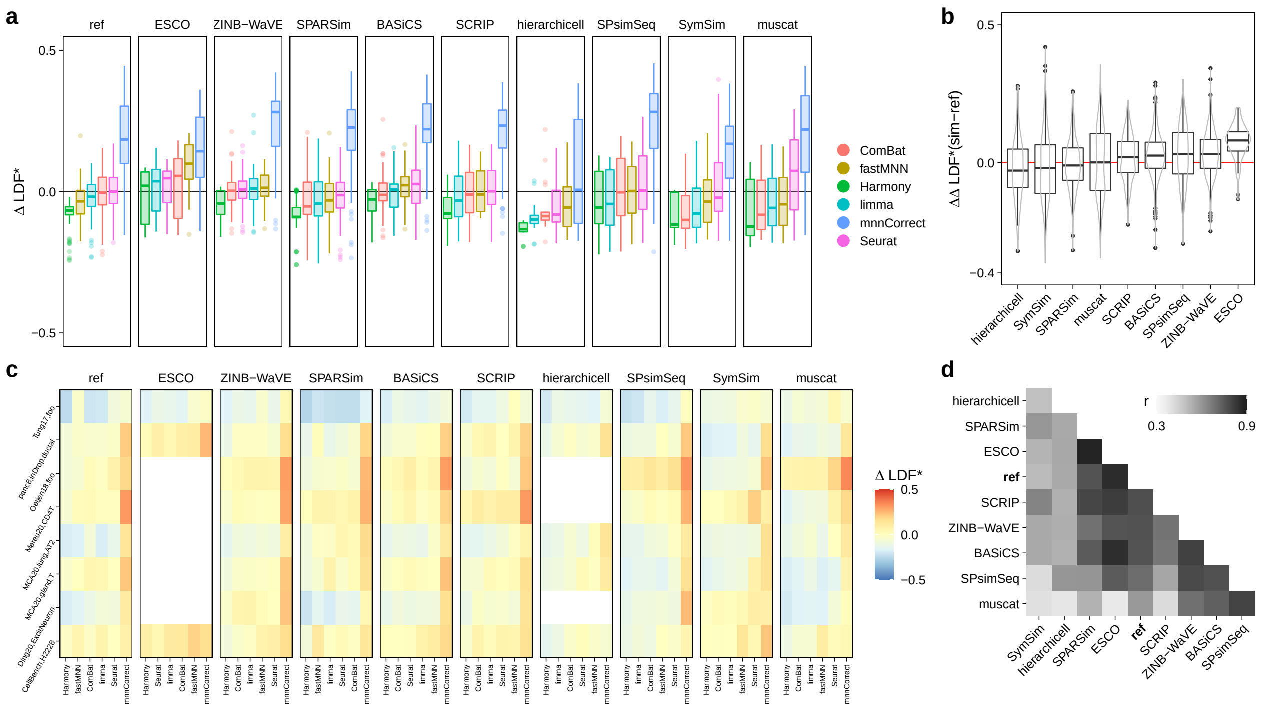

**Supplementary Figure 16:** Comparison of integration results across (experimental) reference and (synthetic) simulated data. (a) Boxplot of batch-level batch correction scores (BCS) across all type $b$ references, simulation and integration methods. (b) Boxplot of difference ($\Delta$) in batch-level BCS obtained from $ref$erence and $sim$ulated data. (c) Heatmap of dataset-level BCS across integration methods (columns) and datasets (rows), stratified by simulator (panels). (d) Heatmap of Spearman's rank correlation ($\rho$) between $\Delta$LDF across datasets and integration methods.

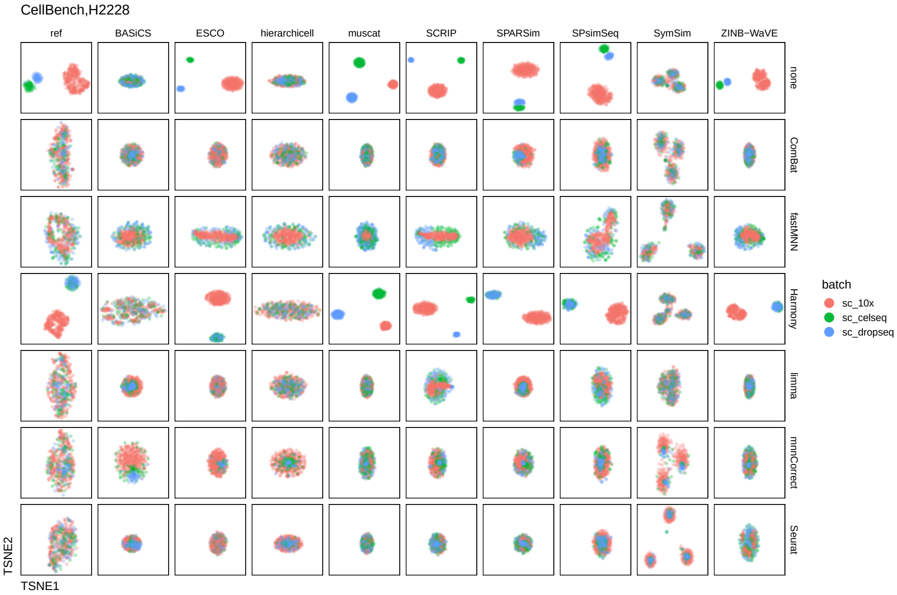

**Supplementary Figure 17:** Comparison of dimension reduction plots across simulators and integration methods for the *CellBench,H2228* dataset; points (cells) are colored by batch.

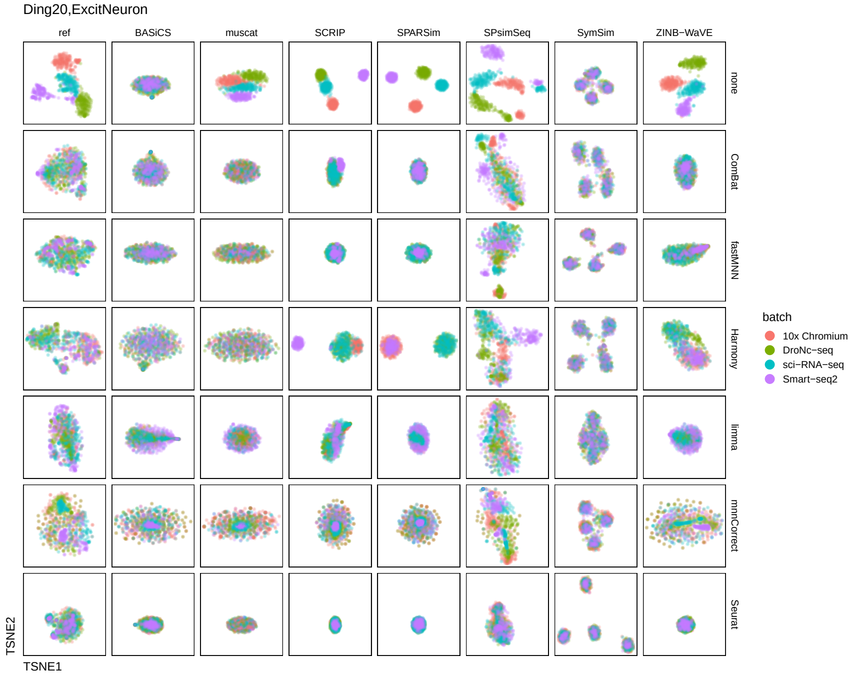

**Supplementary Figure 18:** Comparison of dimension reduction plots across simulators and integration methods for the *Ding20,ExcitNeuron* dataset; points (cells) are colored by batch.

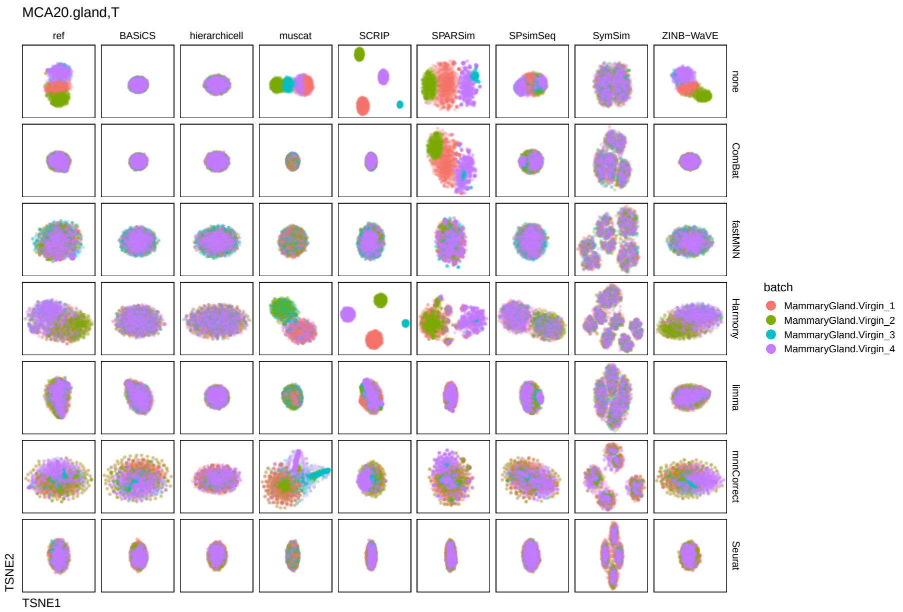

**Supplementary Figure 19:** Comparison of dimension reduction plots across simulators and integration methods for the *MCA20.gland,T* dataset; points (cells) are colored by batch.

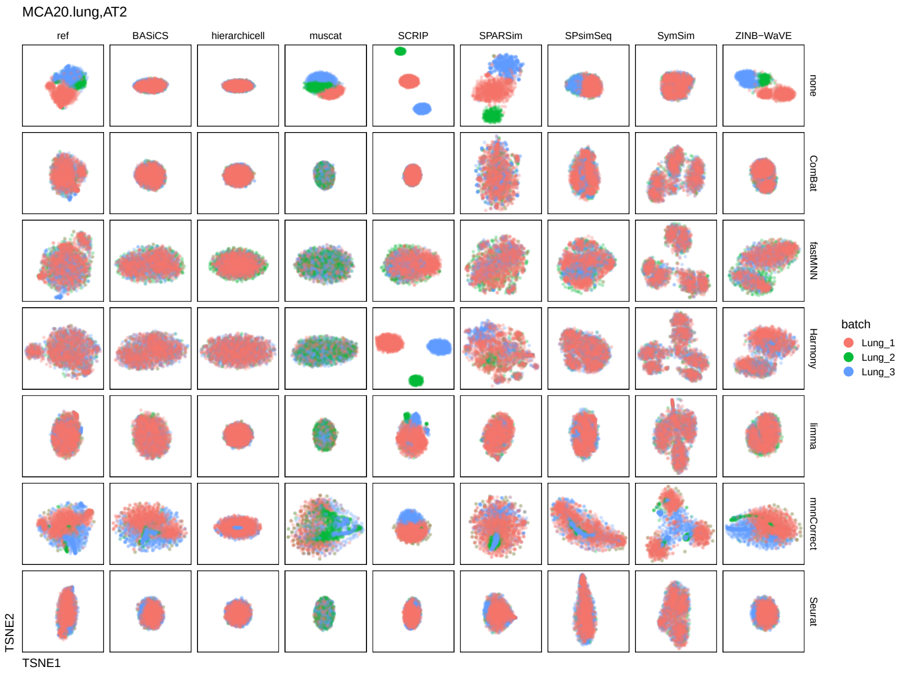

**Supplementary Figure 20:** Comparison of dimension reduction plots across simulators and integration methods for the *MCA20,lung,AT2* dataset; points (cells) are colored by batch.

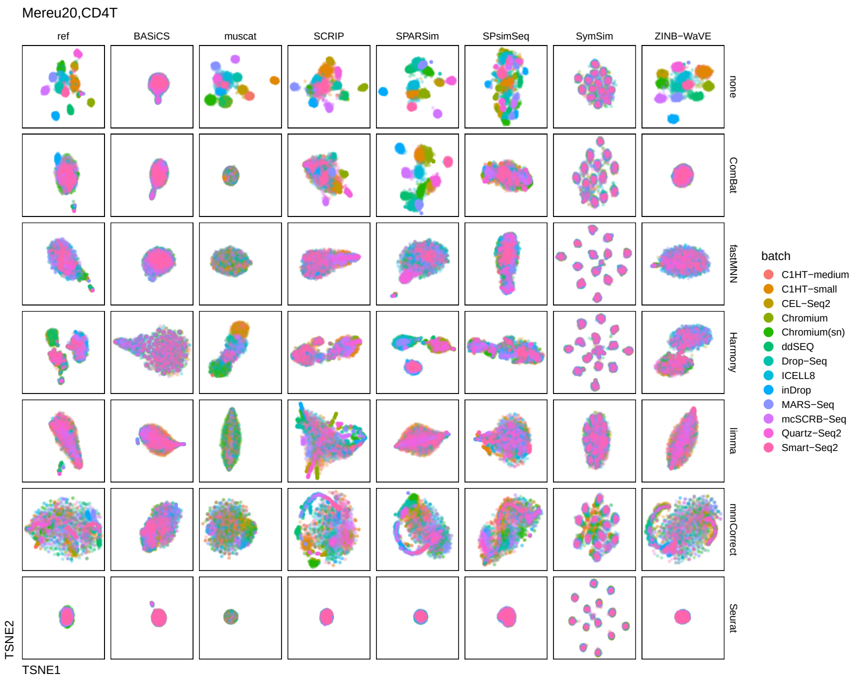

**Supplementary Figure 21:** Comparison of dimension reduction plots across simulators and integration methods for the *Mereu20,CD4T* dataset; points (cells) are colored by batch.

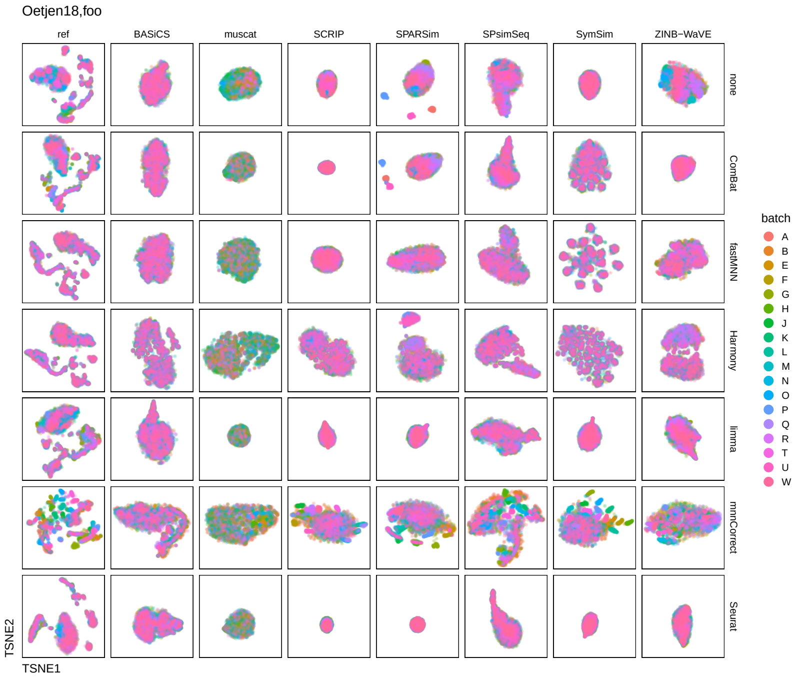

**Supplementary Figure 22:** Comparison of dimension reduction plots across simulators and integration methods for the *Oetjen18* dataset; points (cells) are colored by batch.

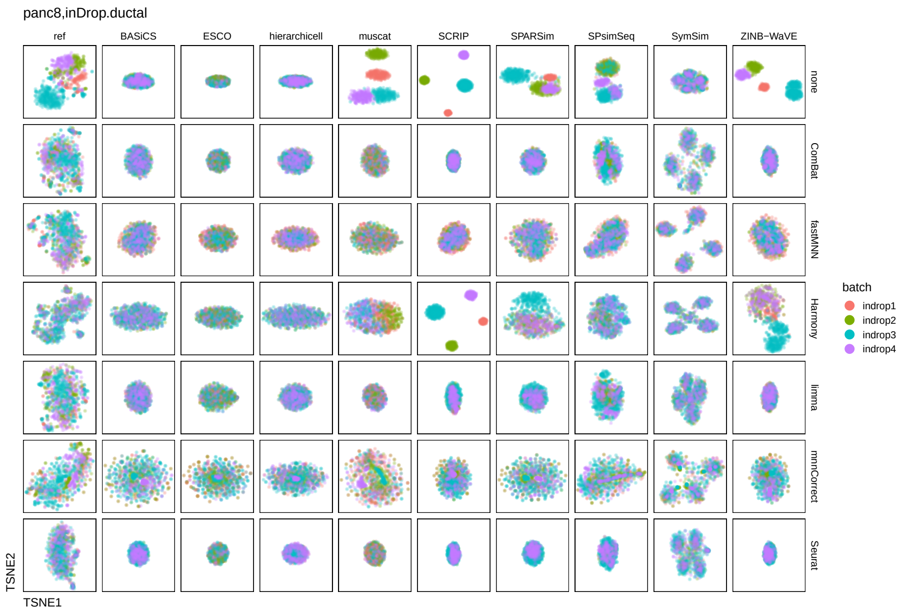

**Supplementary Figure 23:** Comparison of dimension reduction plots across simulators and integration methods for the *panc8,inDrop.ductal* dataset; points (cells) are colored by batch.

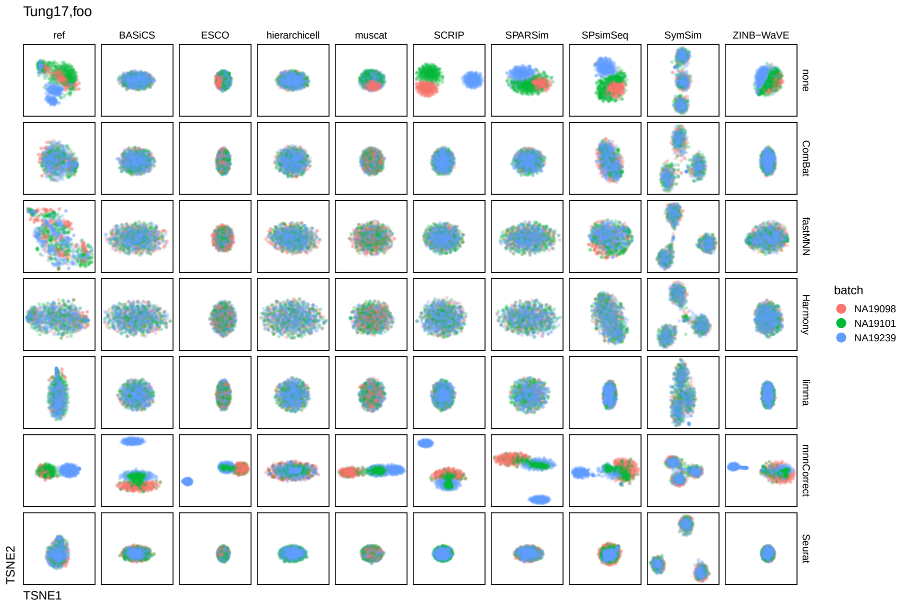

**Supplementary Figure 24:** Comparison of dimension reduction plots across simulators and integration methods for the *Tung17* dataset; points (cells) are colored by batch.

#### 4.2 Summaries

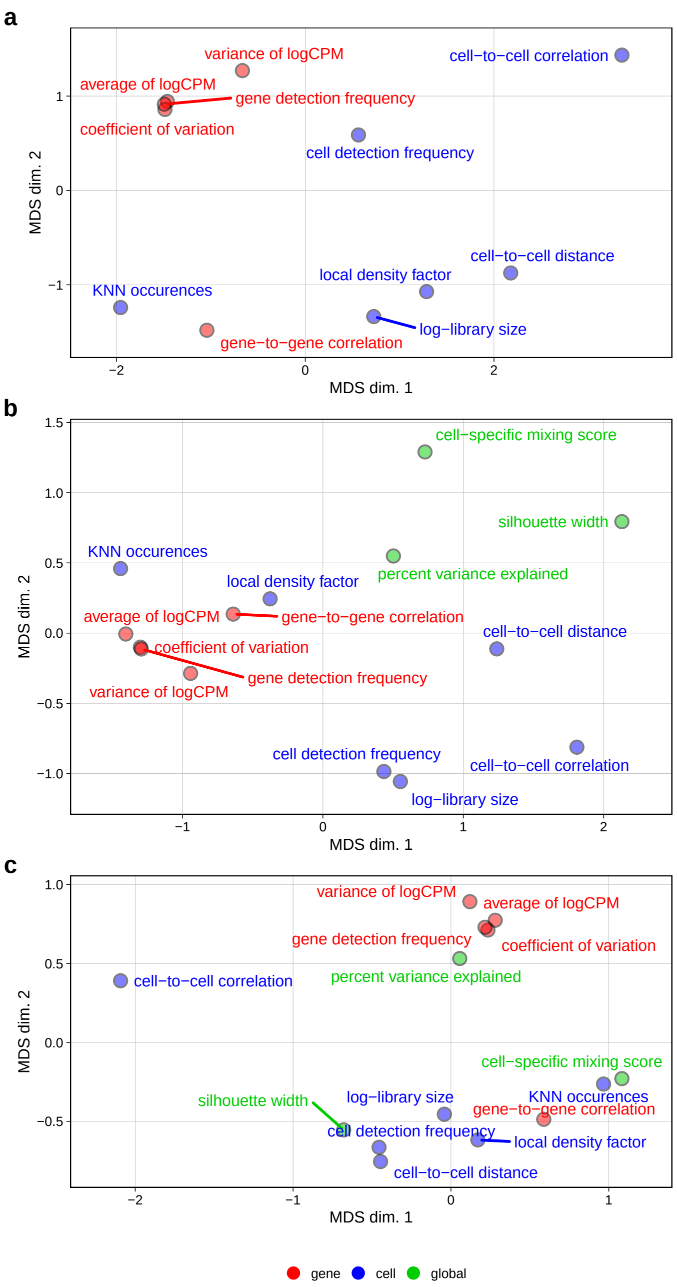

**Supplementary Figure 25:** Multi-dimensional scaling (MDS) plot of per-summary KS statistics across methods and datasets of type $n$ (a), $b$ (b), and $k$ (c). Summaries (points) are colored by their type: gene- (red), cell-level (blue) and, except for type $n$, global (green).

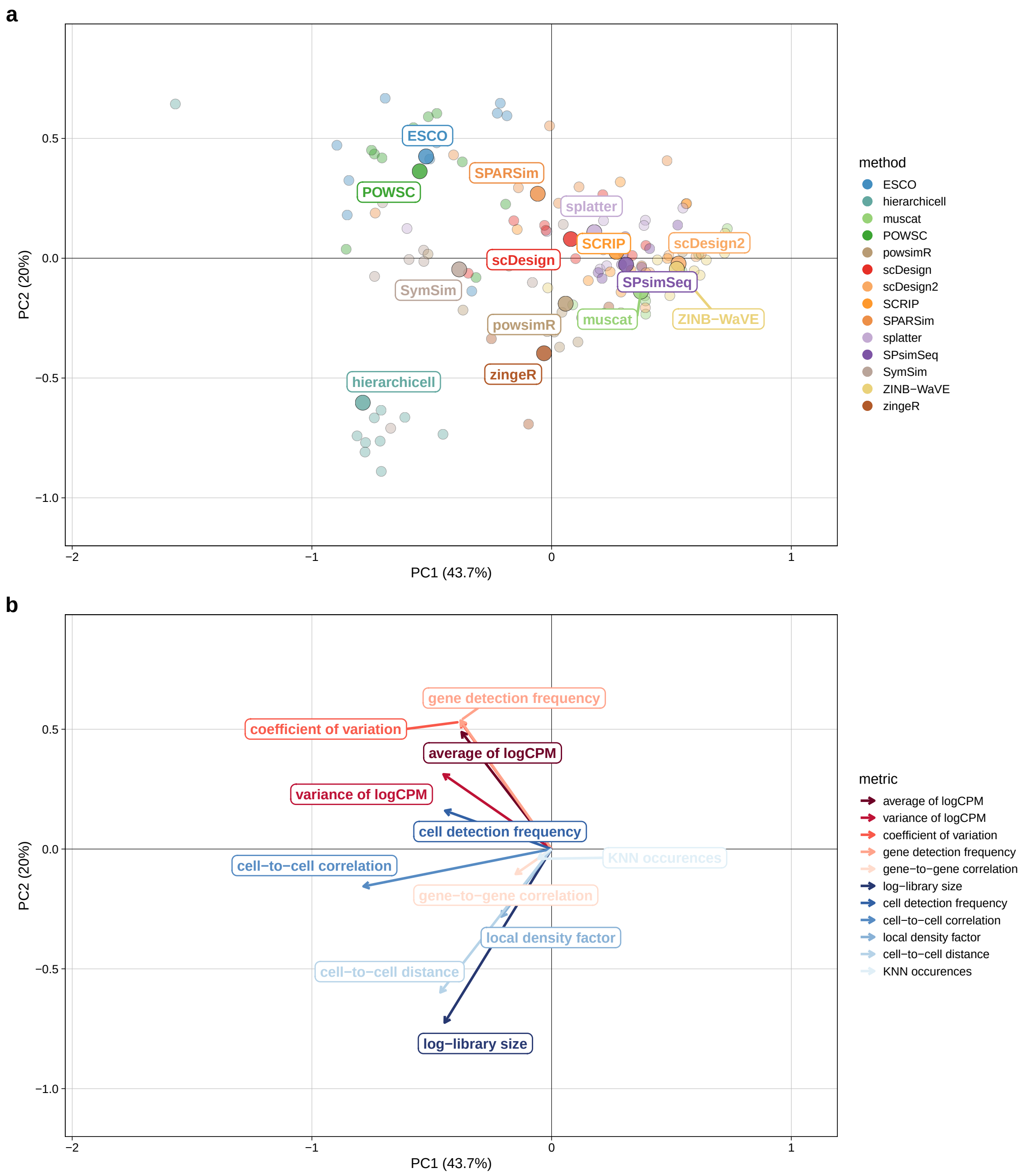

**Supplementary Figure 26:** Principal component (PC) analysis of KS statistics across summaries, methods, and datasets; type $n$. (a) First two PCs. Each small point corresponds to a dataset-method, large points represent per-method averages across datasets. Axis titles indicate the percentage of variance explained by each component. (b) PC loadings. Arrows correspond to summaries and are colored by type (gene- = red, cell-level = blue). (c) Scree plot showing the percentage of variance explained by each PC.

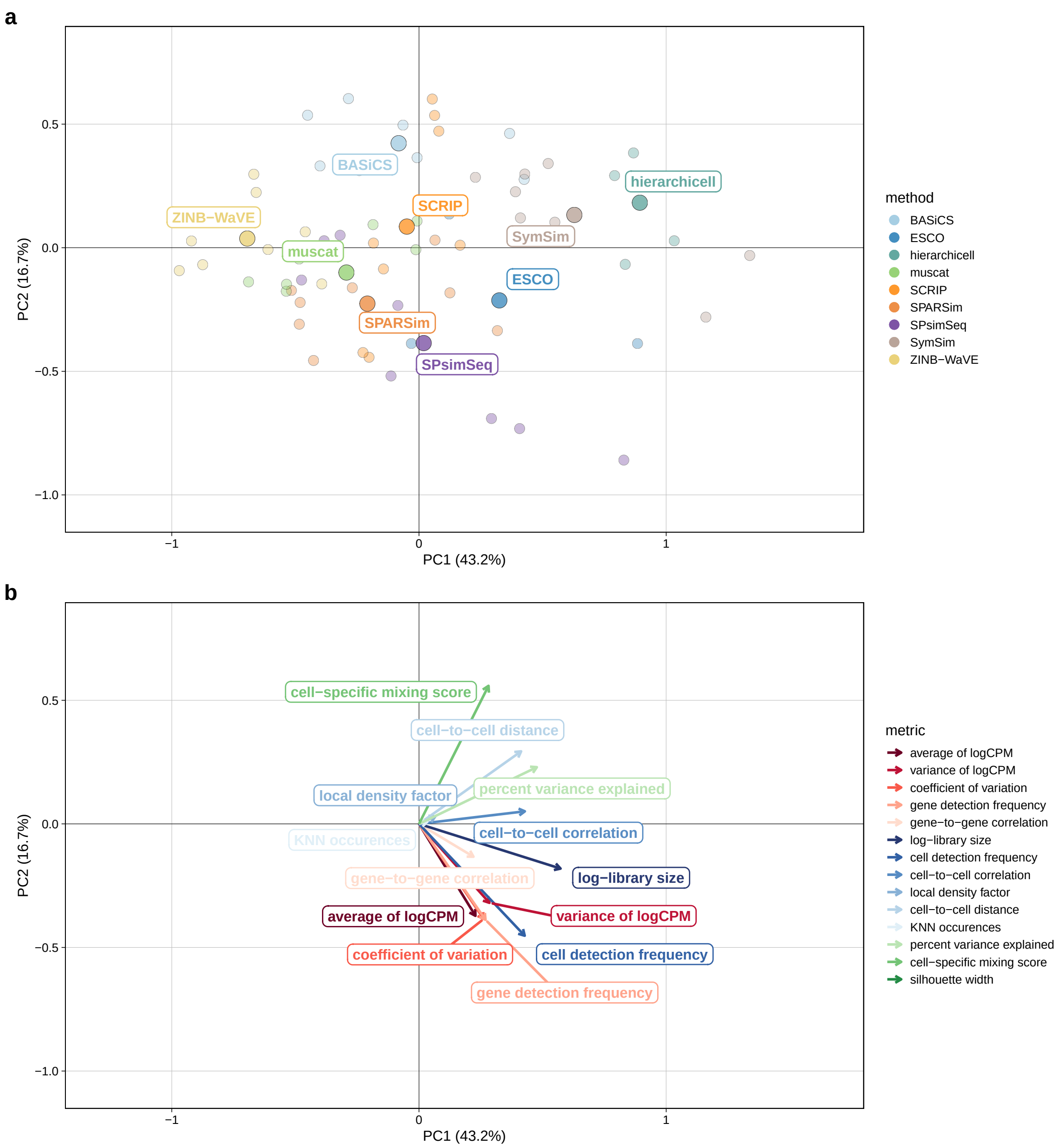

**Supplementary Figure 27:** Principal component (PC) analysis of KS statistics across summaries, methods, and datasets; type *b*. (a) First two PCs. Each small point corresponds to a dataset-method, large points represent per-method averages across datasets. Axis titles indicate the percentage of variance explained by each component. (b) PC loadings. Arrows correspond to summaries and are colored by type (global, gene- or cell-level). (c) Scree plot showing the percentage of variance explained by each PC.

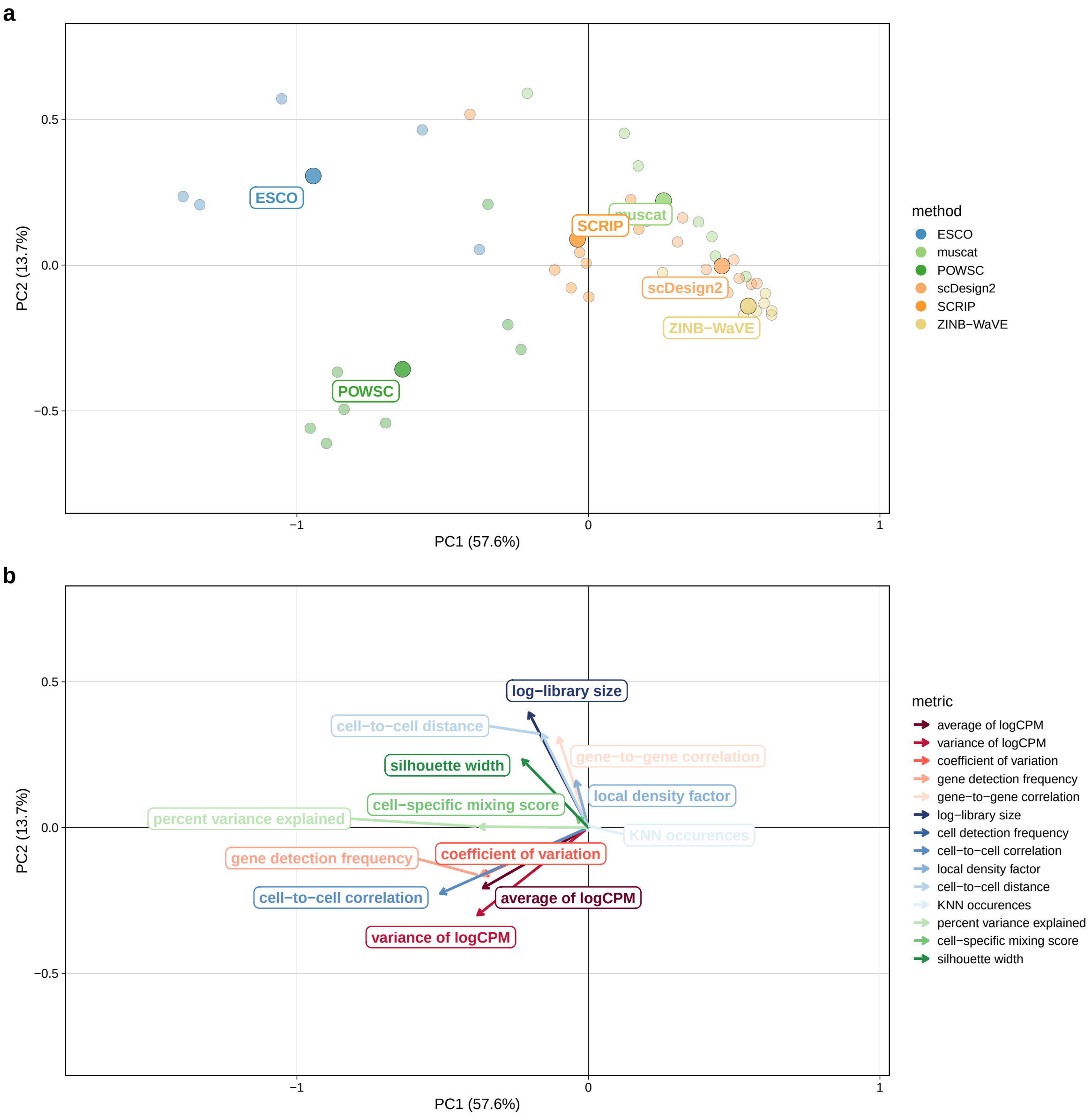

**Supplementary Figure 28:** Principal component (PC) analysis of KS statistics across summaries, methods, and datasets; type $k$. (a) First two PCs. Each small point corresponds to a dataset-method, large points represent per-method averages across datasets. Axis titles indicate the percentage of variance explained by each component. (b) PC loadings. Arrows correspond to summaries and are colored by type (gene- = red, cell-level = blue, global = green). (c) Scree plot showing the percentage of variance explained by each PC.
